# Bioorthogonal Functionalization of Material Surfaces with Bioactive Molecules

**DOI:** 10.1101/2021.10.01.462811

**Authors:** Kern Hast, M. Rhia L. Stone, Zhaojun Jia, Melih Baci, Tushar Aggarwal, Enver Cagri Izgu

**Author notes:** Corresponding author: Izgu, E. C. These authors contributed equally (M. Rhia L. Stone, Zhaojun Jia).

## Abstract

The functionalization of material surfaces with biologically active molecules is crucial for enabling technologies in life sciences, biotechnology, and medicine. However, achieving biocompatibility and bioorthogonality with current synthetic methods remains a challenge. We report herein a novel surface functionalization method that proceeds chemoselectively and without a free transition metal catalyst. In this method, a coating is first formed via the tyrosinase-catalyzed putative polymerization of a tetrazine-containing catecholamine (DOPA-Tet). One or more types of molecule of interest containing *trans*-cyclooctene are then grafted onto the coating via tetrazine ligation. The entire process proceeds under physiological conditions and is suitable for grafting bioactive molecules with diverse functions and structural complexities. Utilizing this method, we functionalized material surfaces with enzymes (alkaline phosphatase, glucose oxidase, horseradish peroxidase), a cyclic peptide (cyclo[Arg-Gly-Asp-D-Phe-Lys], or c(RGDfK)), and an antibiotic (vancomycin). Colorimetric assays confirmed the maintenance of the biocatalytic activities of the grafted enzymes on the surface. We established the mammalian cytocompatibility of the functionalized materials with fibroblasts. Surface functionalization with c(RGDfK) showed improved fibroblast cell adhesion and cytoskeletal organization. Microbiological studies with *Staphylococcus aureus* indicated that surfaces coated using DOPA-Tet inhibit the formation of biofilms. Vancomycin-grafted surfaces additionally display significant inhibition of planktonic *S. aureus* growth.

## INTRODUCTION

Control of surface chemistry is pivotal to the success of many medical devices, which often must operate within biological environments, integrate with host tissues^1, 2^, or fixate biomacromolecules without loss of bioactivity^3, 4^. Because medically useful biomaterials exhibit a wide range of chemical properties, the modification of surface chemical properties is appealing to selectively tailor device-environment interactions or add functionality to the surface while retaining bulk properties and flexibility in material selection. Chemical coating and grafting methods are often used for such purposes, however, for biological applications these methods must be bioorthogonal^5, 6^ and biocompatible^7^. Specifically, the coating must display no toxicity, proceed under physiological conditions, and have high chemoselectivity to maintain the structural integrity of immobilized molecules.

A promising approach to material coating is to employ catechols as molecular building blocks. This approach takes inspiration from the adhesive mussel foot protein Mefp-5^8^, which uses catechol-containing residues to tether mussels to surfaces^9^. Catechols undergo oxidative polymerization, forming supramolecular structures that bind surfaces through bidentate coordination, hydrogen bonding, and π–π stacking^9^. Catecholamines, particularly 3,4-dihydroxy-l-phenylalanine (l-DOPA) and its decarboxylated derivative, dopamine, have been used to produce coatings that can be subsequently modified through Michael addition or Schiff’s base reaction^10^. However, these methods often make use of transition metals to initiate catechol polymerization. These metals are often carcinogenic or cytotoxic^11–13^ or can disrupt the activity of enzymes being grafted onto a surface^14, 15^. The ready coordination of catechols with transition metals^16, 17^ makes complete removal of the metals from coated surfaces challenging. Furthermore, the limited chemoselectivity in post-coating manipulations, e.g., Michael additions and Schiff base reactions, is problematic in the context of biological environments, where biologically active complex molecules, such as nucleic acids, peptides, or proteins, contain multiple nucleophilic amino or thiol groups. Despite major efforts and accumulated knowledge in catechol-based surface chemistry, a systematic surface functionalization approach that is both bioorthogonal and biocompatible remains unreported.

To address these challenges, we report herein a method of chemoselective surface functionalization with complex molecules of interest (MOIs) that proceeds without a free metal catalyst (**Figure 1A**). This method employs tyrosinase to catalyze the putative polymerization of a novel DOPA derivative containing a 1,2,4,5-tetrazine group (DOPA-Tet), which results in the coating of a solid material surface. The coated material can then be grafted with one or more types of MOI that contain *trans*-cyclooctene (TCO) via tetrazine-TCO cycloaddition^18^, here referred to as tetrazine ligation. There are two key innovations in our approach: First, we eliminate free metal catalysts through the enzymatic catalysis of polymerization of the synthetic catecholamine. Catechol polymerization occurs in biology in the production of melanins from l-DOPA—a process called melanogenesis that occurs in the Raper-Mason pathway^19, 20^. In this biosynthetic route, tyrosinase catalyzes the oxidation of l-DOPA and initiates its polymerization. Tyrosinase has been used as a catalyst for the oxidation of aromatic alcohols^21^ and dopamine^22^ but is largely unexplored in its performance with unnatural or more structurally complex DOPA derivatives. Because tyrosinase has been shown to oxidize a wide variety of phenolic and catecholic substrates^21^, we reasoned that it may perform well in promoting the putative polymerization of the DOPA-Tet. Second, the grafting of MOIs to the coated surface is achieved through rapid and highly chemoselective tetrazine ligation. This conjugation chemistry spontaneously occurs in aqueous conditions and produces N_2_ as the sole byproduct^18^. Furthermore, neither the Tet nor TCO groups have been observed in known biology, which avoids the risk of their non-specific covalent interactions within biological media^23^. Synthetic linkers with Tet and TCO groups are readily available and can be incorporated into molecules through standard coupling chemistries. With these features, the tetrazine ligation has an increasing presence in chemical biology^23–26^, and its value has been underscored by its use in technologies that have made it to clinical trials^27^.

**Figure 1.**
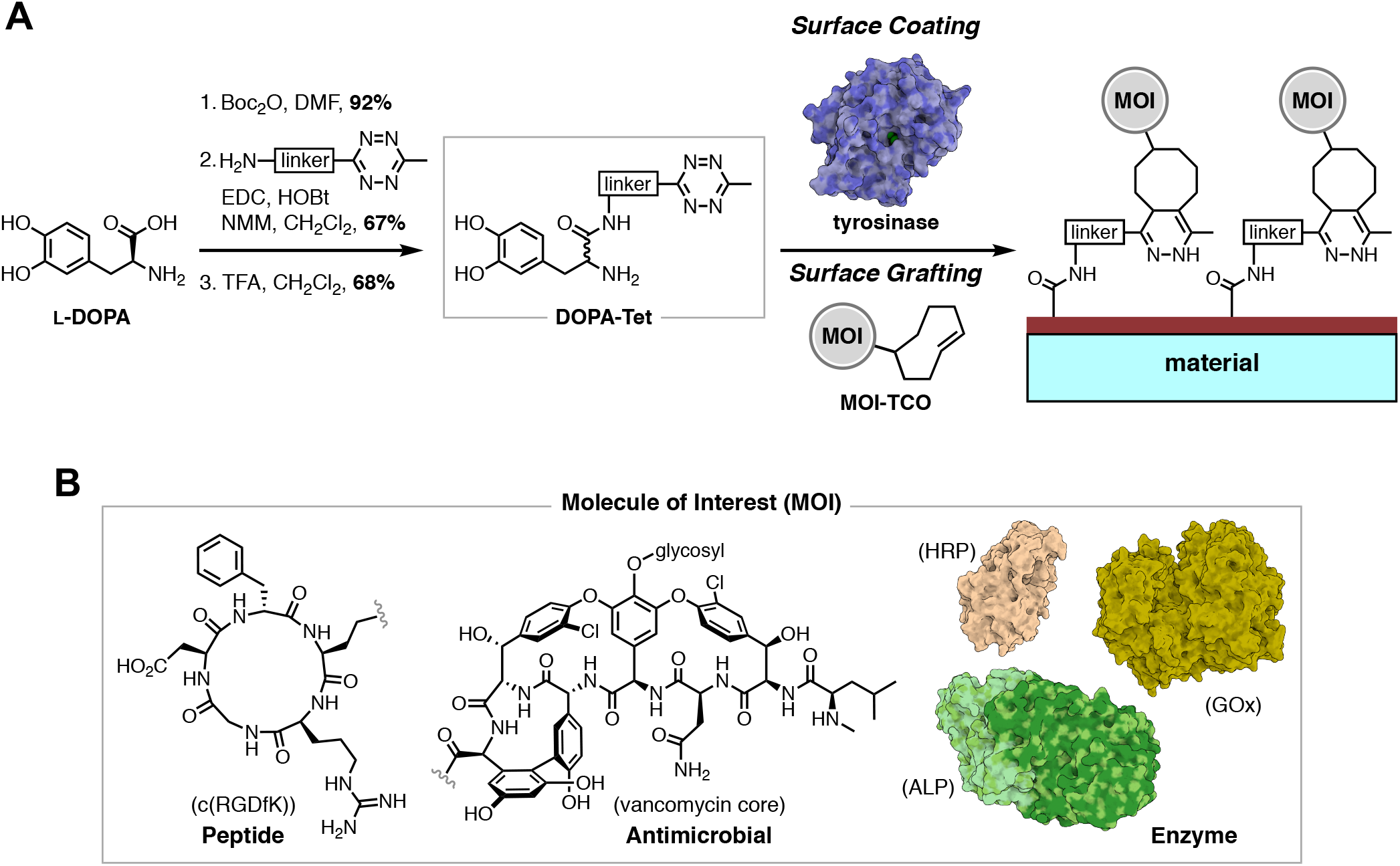
**A**. Overview of the described bioorthogonal material surface functionalization methodology. Linker: – (C_2_H_4_O)_4_–C_6_H_4_–. **B**. The bioactive MOIs grafted on material surfaces in this study. From left to right, these MOIs are depicted in order of increasing molecular size and complexity.

To establish the performance of the described method, we investigated surfaces grafted with molecules that are structurally and functionally distinct (**Figure 1B**). Three enzymes; alkaline phosphatase (ALP), glucose oxidase (GOx), and horseradish peroxidase (HRP); were used to produce surface-immobilized colorimetric assays. ALP is a glycoprotein that catalyzes the hydrolysis of orthophosphate monoesters at alkaline pH^28^. ALP activity is typically measured via the hydrolysis of *para*-nitrophenyl phosphate (p-NPP), which results in a yellow color and an increase in absorbance at 405 nm^29^. GOx is a homodimeric glycoprotein linked via disulfide bonds^30^. Each subunit contains an identical active site that relies on a tightly, noncovalently bound FAD cofactor and oxidizes glucose to gluconolactone, producing hydrogen peroxide (H_2_O_2_) as a byproduct^30^. HRP is a family of isoenzymes that are composed of a single polypeptide with an iron(III) protoporphyrin IX (commonly referred to as a “heme group”) cofactor^31^. HRP uses H_2_O_2_ to oxidize a variety of organic substrates, including several small-molecule chromophores that undergo a color change when oxidized^32^. In this study, we used 2,2’-azino-bis(3-ethylbenzthiazoline-6-sulfonic acid) (ABTS), whose oxidation by HRP produces a blue-green color and an increase in absorbance at 405 nm. When GOx and HRP are colocalized, the H_2_O_2_ produced by GOx can be utilized by HRP to produce a colorimetric signal.

Additionally, we investigated the effect on cell adhesion and viability of grafting surfaces with the cyclic peptide cyclo[Arg-Gly-Asp-D-Phe-Lys] (c(RGDfK)) and vancomycin. c(RGDfK) includes the common Arg-Gly-Asp motif that binds adhesive proteins on cell surfaces, most notably integrins, and has been shown to improve cell adhesion to surfaces^33^. We cultured fibroblasts (NIH3T3) on c(RGDfK)-grafted discs of NanoECM™, which is a randomly oriented, electrospun polycaprolactone product designed to imitate decellularized tissue. Cell viability, adhesion, and morphology were examined with confocal laser scanning microscopy (CLSM), scanning electron microscopy (SEM), and 3-(4,5-dimethylthiazol-2-yl)-2,5-diphenyl tetrazolium bromide (MTT) cell proliferation assays. Vancomycin is a glycopeptide antibiotic effective against several clinically relevant Gram-positive bacteria, including *Staphylococcus aureus*^34^. *S. aureus* was cultured on vancomycin-grafted surfaces and the effect on cell viability and growth was studied with growth inhibition assays and live cell microscopy.

## RESULTS AND DISCUSSION

### Tetrazine-Modified DOPA (DOPA-Tet), a Synthetic Derivative of l-DOPA, Can Be a Substrate for Tyrosinase

We developed a scalable synthetic route for DOPA-Tet using readily available l-DOPA and (3-phenyl-6-methyl-1,2,4,5-tetrazine)-PEG4-amine (here referred to as Tet-PEG4-NH_2_) (**Figure 1A**). Briefly, l-DOPA was *N*-Boc protected using Boc_2_O and Tet-PEG4-NH_2_ was installed via EDC/HOBt coupling. Subsequent removal of Boc in a mixture of TFA/CH_2_Cl_2_ provided DOPA-Tet in 42% yield over 3 steps (see SI for details). The amide bond formation was facilitated with *N*-methylmorpholine (NMM), which resulted in racemization as determined by optical rotation analysis. We believe this is likely inconsequential in the context of our coating approach, as tyrosinase is known to use both l- and D-enantiomers of DOPA as a substrate^35^. l-DOPA is highly prone to degradation under basic and oxidative conditions^36^, rationalizing the protection of both the amino and hydroxyl groups prior to chemical modifications on the catecholamine structure, which we previously demonstrated to be effective^37^. In the current study, however, we observed that the structural integrity of the catecholamine during the tetrazine installation was maintained without hydroxyl protection. Omitting this step allowed for a more straightforward protecting group chemistry and improved the overall step economy.

Central to this coating method is the use of tyrosinase to putatively catalyze the polymerization of DOPA-Tet, leading to its deposition on the surface. Because DOPA-Tet is not a natural substrate of tyrosinase, we evaluated tyrosinase activity on DOPA-Tet by coating two types of materials, polypropylene and polyethylene terephthalate (PET), with either l-DOPA or DOPA-Tet. For each surface, we evaluated coating either with or without tyrosinase (**Figure 2A**). For both catecholamines, the tyrosinase-containing samples displayed a darker color after 1 hour compared to the initial reaction solutions, whereas the samples without tyrosinase did not change color. When washed with deionized water, both tyrosine-containing samples left clearly visible coatings on the material surface. In contrast, no coating was visible from the l-DOPA sample without tyrosinase, and only a small amount of coating was visible from the DOPA-Tet sample without tyrosinase.

**Figure 2.**
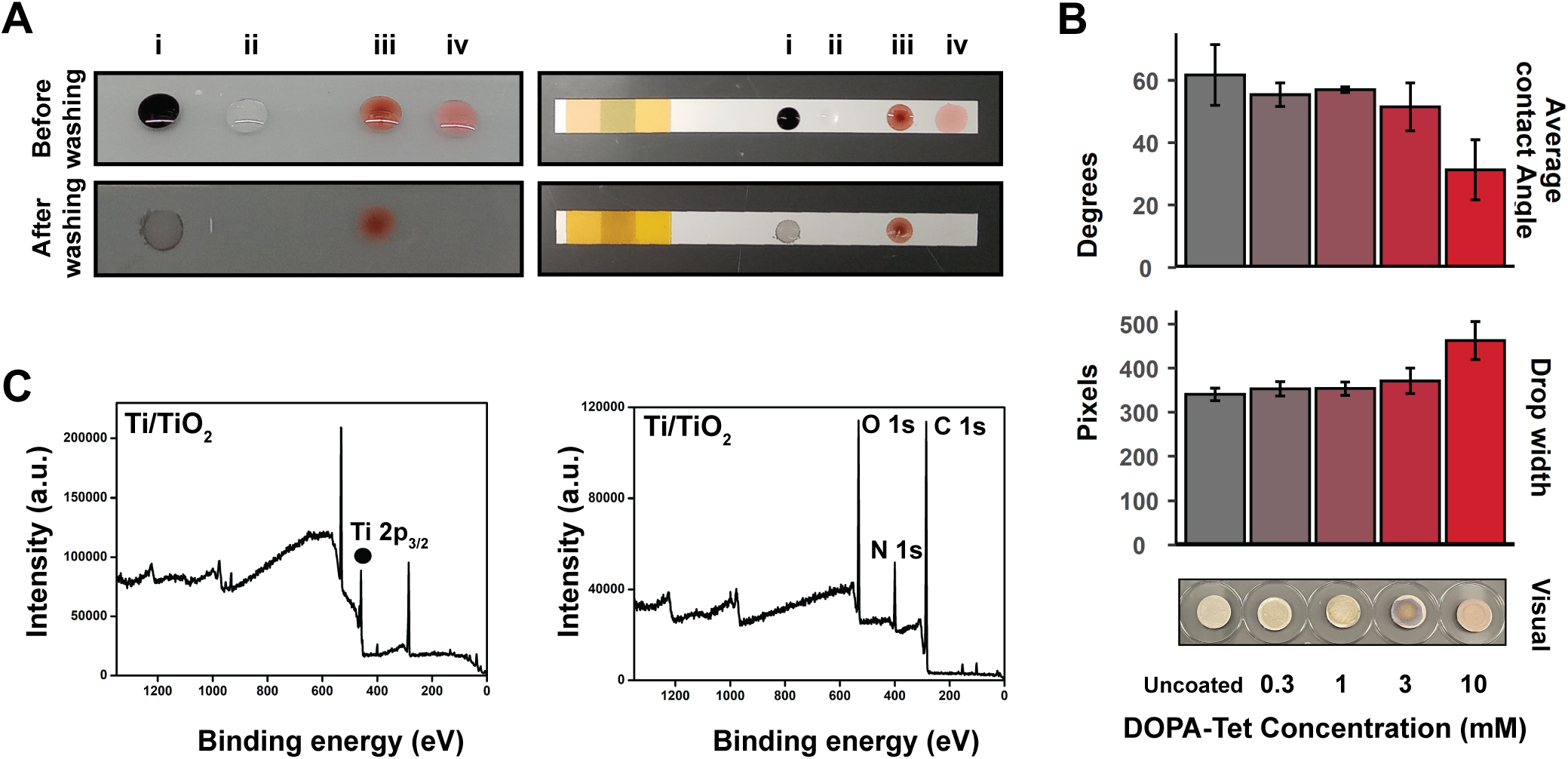
**A**. (Left) Polypropylene and (Right) polyethylene terephthalate surfaces incubated with (i) l-DOPA and tyrosinase, (ii) l-DOPA, (iii) DOPA-Tet and tyrosinase, and (iv) DOPA-Tet. Samples are shown (Top) before and (Bottom) after washing away the coating solutions. **B**. (Top, Middle) Contact angle measurements using sessile drop method on coated and uncoated Ti discs. (Bottom) DOPA-Tet coated and uncoated Ti discs. Error bars denote 1 standard deviation. **C**. XPS spectra of Ti/TiO_2_ (Left) before and (Right) after coating with 10 mM DOPA-Tet. The black dot denotes the Ti substrate peak.

### Assessment of DOPA-Tet Coatings

Contact angles for water on both uncoated and coated Ti, Si, and glass were collected using the sessile drop method and analyzed (**Figure 2B** and **Figure S1**). Before coating, glass was found to be much more hydrophilic than Ti or Si, with contact angles of 19.0°, 61.8°, and 71.6°, respectively. However, irrespective of the material, the coated surfaces showed similar hydrophilicity and a trend of decreasing contact angle with increasing concentration of DOPA-Tet. Specifically, 0.3 mM coated contact angles were 55.5°, 67.8°, and 45.5°; and 10 mM coated contact angles were 31.3°, 42.8°, and 30.0° for Ti, Si, and glass, respectively. These results support the efficient coating of the surface even at relatively low concentrations of DOPA-Tet.

Coated Ti, Si, and glass were investigated with X-ray photoelectron spectroscopy (XPS) (**Figure 2C**) and attenuated total reflectance Fourier-transform infrared spectroscopy (ATR-FTIR) (**Figure S2**). In XPS, the material substrate peaks were observed in the uncoated samples but were greatly diminished in the coated samples, being replaced with the C, N, and O peaks of organic species. This replacement of the substrate signal indicated that organic materials from DOPA-Tet were deposited on the surface. ATR-FTIR was utilized to observe coating materials both before and after the application of tyrosinase. DOPA-Tet both before and after incubation with tyrosinase was characterized as deposits from solutions added directly onto the FTIR diamond sensor. These spectra were compared to references for Tet-PEG4-NH_2_, which was characterized by us using the same approach, and a reported spectrum for l-DOPA^38^. We observed peaks in the 2800–3800 cm^−1^ region representing the stretching vibration of O–H (catechol) and N–H (either RNH_2_ or R_2_NH). The strength of the C=O stretching peak at 1670 cm^−1^, corresponding to the amide bond, was reduced in the tyrosinase-treated DOPA-Tet sample relative to the untreated sample. These observations are consistent with those for the coatings obtained with a previously reported amide-containing DOPA analog^37^. The C=C and N–N stretching vibration peaks within the region of 500–1600 cm^− 1^ observed in Tet-PEG4-NH_2_ were reflected in the coated sample, supporting the existence of tetrazine residues on the coating.

In addition to these instrumental analyses conducted directly with the coatings, we sought to investigate whether the tyrosinase-mediated surface deposition mechanism is affected by the amino group substitution of the parent DOPA structure. Notably, *N*-Boc protection inhibited the putative polymerization of DOPA-Tet when tyrosinase was added (**Figure S3**), suggesting that accessibility of the free amino group is a necessary step for supramolecule formation. This is in agreement with observations made for the polymerization of a related amido-containing catecholamine^37^ and dopamine^39^.

Finally, we assessed the stability of the coatings in different media. We incubated coated glass surfaces for 5 days at 37 °C in buffers with pH 4.5 to 9.5 and in 10% dimethylsulfoxide (DMSO) in PBS (pH 7.4) (**Figure S4**). No degradation of the coating was observed in the samples incubated in pH 4.5 to 8.5. However, delamination and degradation of the coating was observed in the sample incubated at pH 9.5, and the sample subjected to 10% DMSO in PBS was partially solubilized. These observations suggest that the coating is stable in near neutral and slightly acidic conditions.

### *Trans*-Cyclooctene-Containing Molecules Undergo Facile, Bioorthogonal Ligation with Coated Surfaces

TCO-containing MOIs were obtained commercially or prepared using protocols appropriate for each MOI. Briefly, Cy5-TCO was purchased and used as received. c(RGDfK)-TCO was obtained from c(RGDfK) and TCO-PEG8-NHS. Vancomycin-TCO was synthesized from vancomycin-HCl and TCO-PEG8-amine. Building on previous works that allow peripheral vancomycin modifications^40, 41^, we attached the TCO handle at the C-terminal carboxylic acid of vancomycin by an amide bond coupling reaction and without the use of protecting groups. ALP, GOx, and HRP, enzymes commonly used in colorimetric assays (**Figure 3A**), were modified with TCO using TCO-PEG24-COOH under EDC/NHS coupling conditions. When preparing the TCO-conjugated enzymes, excess amounts of TCO-PEG24-COOH were used to attain efficient reaction kinetics, which likely resulted in the conjugation of multiple TCO groups onto each enzyme. Examination of ALP, GOx, and HRP crystal structures (PDB IDs 3TG0^42^, 3QVP^43^, and 1H58^44^, respectively) showed that the enzyme active sites did not contain lysine residues, and thus lysine-TCO conjugations were unlikely to impact enzyme activity. To verify that the TCO-PEG24 attachment did not significantly disrupt enzyme activity, we first conducted assays with solutions of ALP-TCO and p-NPP in Tris-buffered saline (TBS) at pH 8.5 or GOx-TCO, HRP-TCO, D-glucose, and ABTS in phosphate buffered saline (PBS) at pH 7.4. We similarly prepared solutions wherein one of the assay components was missing. The complete assays produced the expected color changes, but any solution which omitted an assay component produced no color change (**Figure S5**). We further compared the kinetics of the native and TCO-conjugated enzymes using UV-Vis spectrophotometry. Assays were produced as described from either TCO-conjugated or native enzymes. Estimations of Michaelis-Menten parameters using non-linear least squares regression showed that the native and TCO-conjugated systems showed similar activities (**Figure 3B**). ALP had a Michaelis constant of *K*_*m*_ = 0.207 mM and ALP-TCO showed *K*_*m*_ = 0.151 mM (p = 0.162). A similar relation was observed for the GOx/HRP system, where the native enzymes and the TCO-conjugated enzymes showed similar *K*_*m*_ (13.5 and 9.89 mM, respectively; p = 0.153).

**Figure 3.**
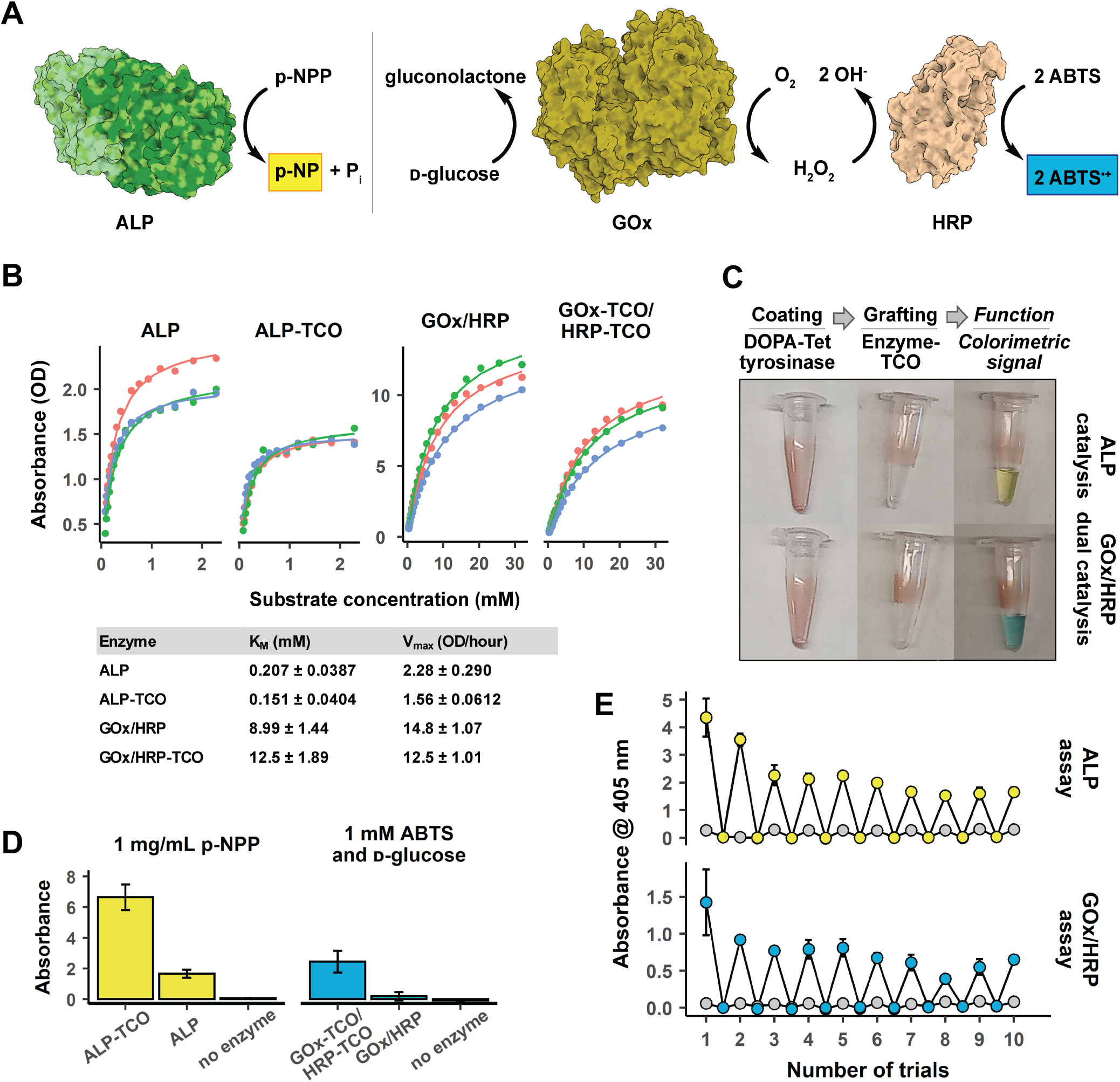
Activities of TCO-conjugated ALP, GOx, and HRP **A**. Schematic representations of (Left) ALP and (Right) GOx/HRP activities. **B**. (Top) Michaelis-Menten Plots and (Bottom) Michaelis-Menten parameters for native and TCO-containing ALP and GOx/HRP. **C**. Images of visual changes after (Left) coating, (Middle) grafting, and (Right) incubation with substrate solutions for (Top) ALP and (Bottom) GOx/HRP. **D**. Absorbance change of substrate solutions incubated in microcentrifuge tubes coated/grafted with TCO-conjugated enzymes, native enzymes, or no enzymes. Error bars denote 1 standard deviation. **E**. Absorbance outputs after repeated uses of (Left) ALP and (Right) GOx/HRP immobilized assays. Assay values are shown in color, negative controls shown in gray.

The preservation of activity in the TCO-attached molecules suggests that little enzyme degradation or denaturation occurred during the conjugation process and the TCO linker did not significantly inhibit the enzymes. These results also confirm that both GOx-TCO and HRP-TCO must be present to produce a color change in an ABTS/glucose solution and that the coenzymes of GOx and HRP, FAD and heme, respectively, are not lost or disrupted by the TCO modification.

The second step of the surface functionalization method, grafting MOIs to the coated surface, relies on the ligation of TCO-containing MOIs to the tetrazine-decorated coating. To verify tetrazine ligation, both uncoated and DOPA-Tet-coated titanium discs were incubated with either sulfo-Cy5 amine (Cy5) or Cy5-TCO (**Figure S6**). Titanium was chosen to minimize the non-specific binding of the fluorophore to the surface, which was observed when using plastic surfaces. All samples showed similar fluorescence intensity before rinsing, however after rinsing with methanol, the sample that was both coated and incubated with Cy5-TCO exhibited significantly higher fluorescence intensity than the uncoated samples or the samples incubated with Cy5. This result indicates that the proposed tetrazine-TCO ligation was successful in retaining an organic small-molecule on a solid surface, which motivated us to expand this methodology for decorating materials with structurally complex and biologically active MOIs.

The general process for functionalizing a surface with an MOI was as follows. The surface was first incubated with a solution of 1–10 mM DOPA-Tet and 2,500 U/mL tyrosinase in PBS for at room temperature for 1–2 hours. After incubation the surface was washed with water or PBS, after which a red-colored film persisted on the surface. This film was observable with concentrations as low as 1 mM DOPA-Tet and serves as a visual indicator of successful coating. When coating with concentrations significantly above 10 mM DOPA-Tet, excessive aggregation and uneven coating were observed. It should also be noted that, due to our synthetic route, a solution of DOPA-Tet will be acidic. Acidified coating solutions from high concentrations of DOPA-Tet were observed to inhibit tyrosinase activity and polymerization. If coating with concentrations above 10 mM, the pH of the coating may need to be adjusted or adequately buffered. After coating and washing away of residual coating material, a solution of 0.1–0.5 mM TCO-conjugated MOI in PBS was incubated on the coated surface at room temperature for 1–2 hours, after which residual solution was washed from the surface. The choice of wash solution was found to be important in removal of unreacted MOI from the surface. We observed that Cy5 and Cy5-TCO were not readily removed from most surfaces with aqueous solutions, and methanol proved more effective. We also observed that ALP was more readily washed from coated microcentrifuge tubes with ionic buffers than with water (See SI for details, **Figure S7**). After ligation, the coating color diminishes and may become less visible. Checking for differences in surface hydrophobicity with a droplet of water proved useful in verifying that the coating was still present.

### Grafted Enzymes Retain Their Activity to Produce Surface-Immobilized Colorimetric Assays

To investigate the efficacy of grafting complex protein catalysts, we prepared surface-immobilized colorimetric assays using two different enzyme systems: ALP and a combination of GOx and HRP. For the ALP-based assay, Eppendorf LoBind® microcentrifuge tubes were coated with DOPA-Tet and incubated with either ALP-TCO, native ALP, or no enzyme, and then washed with PBS (**Figure 3C**). The catalytic activities on a solution of 1 mg/mL p-NPP over 1 hour were assessed via measurements of their absorbance at 405 nm (**Figure 3D**). The surfaces treated with ALP-TCO showed significantly more activity (*A* = 6.65) than those treated with ALP (*A* = 1.66, p = 0.005) or untreated (*A* = 0.05, p = 0.005). Native ALP is not able to covalently bond with the tetrazine functionalized surface, and as such any activity exhibited by the ALP-treated surface is likely due to ALP that persists after washing via non-specific interactions.

We also prepared an assay utilizing GOx and HRP in combination. Microcentrifuge tubes were coated DOPA-Tet and incubated with either GOx-TCO and HRP-TCO, GOx and HRP, or no enzymes, and then washed with PBS. Their activities on a solution of 1 mM D-glucose and 1 mM ABTS over 1 hour were assessed via absorbance measurements (**Figure 3D**). The GOx-TCO/HRP-TCO treated surfaces exhibited significantly more activity (2.45 OD) than either the GOx/HRP treated (0.187 OD, p = 0.0003) or coated-only (−0.07 OD, p = 0.004) surfaces. As with ALP, it is expected that the native enzymes are removed during washing, and some activity is observed in those samples due to non-specific binding to the tube.

It is notable that much greater activity was observed from the native ALP samples than from the native GOx/HRP samples. This observation could have multiple explanations. Both GOx and HRP must be present to successfully convert the substrate, and the process will be limited if the concentration of either enzyme is reduced. Also, HRP is significantly smaller than ALP (44 kDa and 160 kDa, respectively), and GOx and HRP have fewer charged surface residues than ALP. Differences in enzyme size and surface chemistry could affect the solubility and degree of interaction with the coated surface^45^, and consequently the efficiency of its removal during washing.

### Grafted Surfaces Are Stable in Biological Media and After Repeated Use

We assessed the stability of surface-grafted enzymes when the surface was subjected to repeated assay and wash conditions (Figure 3E). Microcentrifuge tubes coated and grafted with either ALP or a combination of GOx and HRP were repeatedly incubated with substrate and buffer solutions with wash steps in between. Absorbance measurements at 405 nm showed that the tubes showed little loss of activity over 10 cycles.

We also assessed the stability of grafted MOIs under biological conditions by incubating in human serum (**Figure S8**). Two vials, one coated and one coated and grafted with ALP-TCO, were incubated in human serum for 5 days at 37 °C and then washed with PBS. Subsequent incubation with 1 mg/mL p-NPP showed ALP activity in the grafted vial, but no activity in the vial that was only coated. These results indicate that the ALP-TCO grafted surface was stable in human serum for prolonged periods. Moreover, these results also show that the native ALP present in serum^28^ had negligible binding to the coating, supporting the bioorthogonality of the approach.

### Functionalization of NanoECM with c(RGDfK) Improves Fibroblast Adhesion and Cytoskeletal Organization

We evaluated the cytocompatibility of c(RGDfK)-grafted NanoECM with fibroblasts (NIH3T3) using an MTT assay of material extracts. NanoECM samples were either untreated or coated/grafted using DOPA-Tet/c(RGDfK)-TCO. Extracts of the samples were prepared in DMEM and fibroblasts seeded in well plates were cultured in these extracts for 1 or 3 days. The metabolic activity of cells cultured in extracts of coated/grafted NanoECM was comparable to that of cells cultured in extracts of uncoated NanoECM and negative controls (DMEM), with day 3 absorbances being more than double those of day 1 (**Figure S9**). Positive controls (10% DMSO-containing DMEM) at both days showed much lower activity than all other groups. These results show that cell proliferation was unaffected by the coating and c(RGDfK) grafting, suggesting that the coating exhibits minimal mammalian cytotoxicity.

Next, we assessed the effect of NanoECM on fibroblast morphology through CLSM and SEM imaging. Both the untreated and c(RGDfK)-grafted NanoECM samples were seeded and cultured with fibroblasts. For confocal imaging, the cells were fixed after 3 hours and stained with a FAK100 kit. Nuclei (via DAPI), vinculin (via antivinculin), and F-actin (via TRITC-phalloidin) were stained blue, green, and red, respectively (**Figure 4A**). The cells on the coated samples showed superior organization of cytoskeletal components vinculin and F-actin. For SEM analyses, the cells were fixed at different time points ranging from 1 hour to 3 days. These samples were gold sputtered prior to SEM imaging (**Figure 4B**). Similar to the results of the CLSM study, we observed greater spreading and more extended projections of cells cultured on c(RGDfK)-grafted NanoECM samples compared to those cultured on untreated samples. This effect was more pronounced at short time scales (up to 6 hours), after which both samples showed similar cell morphologies. The observations of these two studies suggest superior fibroblast adhesion to the NanoECM surfaces that had been coated/grafted with DOPA-Tet/c(RGDfK)-TCO.

**Figure 4.**
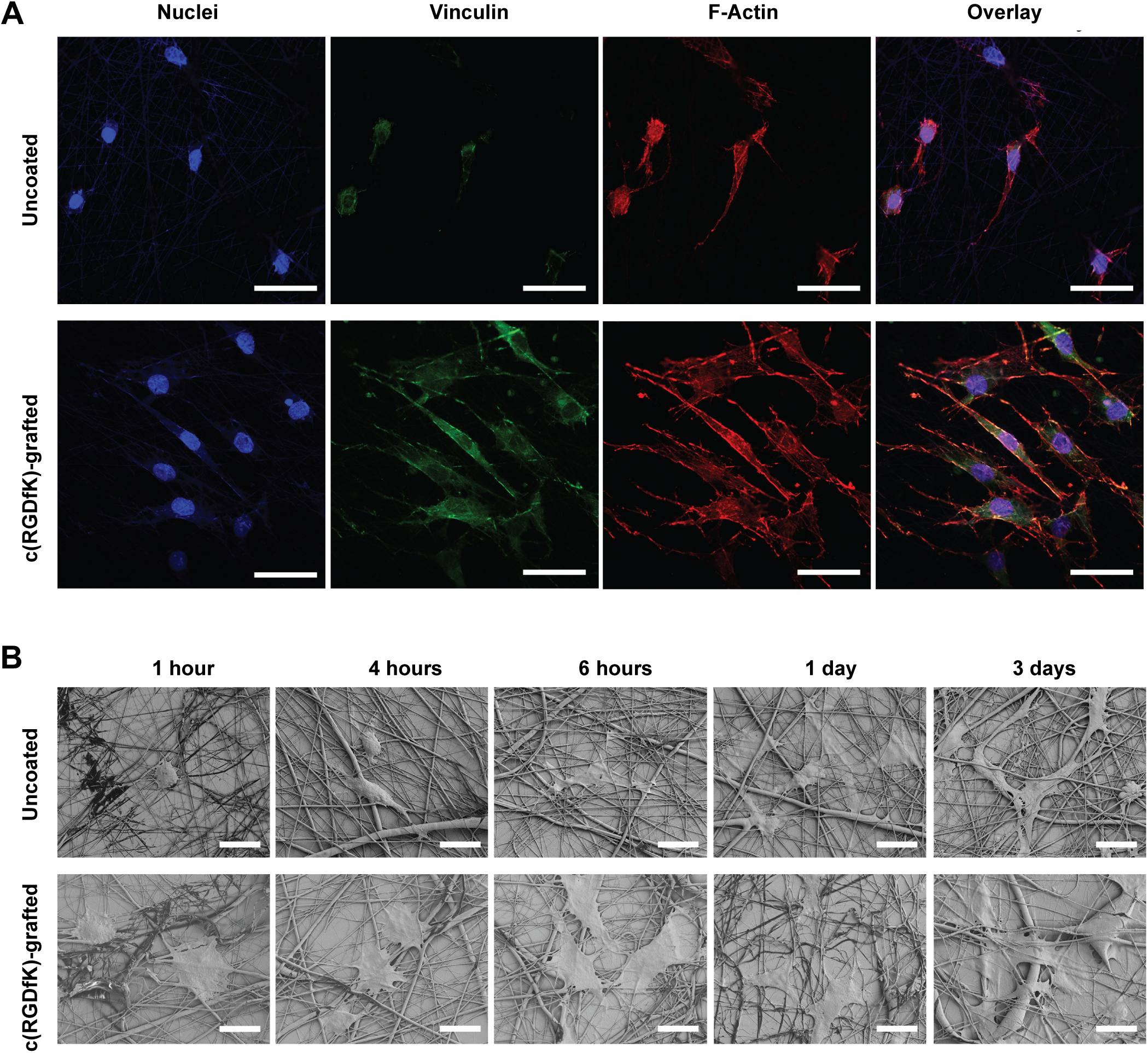
Cytocompatibility studies of NIH3T3 fibroblasts. **A**. Confocal images of cell adhesion and cytoskeleton organization on uncoated and c(RGDfK)-grafted NanoECM. Nuclei (via DAPI), vinculin (via antivinculin), and F-actin (via TRITC-phalloidin) are stained blue, green, and red, respectively.

### Grafted Vancomycin Inhibits *S. Aureus* Growth and Biofilm Formation

We investigated the microbial inhibitory effect of surfaces coated/grafted using DOPA-Tet/vancomycin-TCO. We first confirmed that vancomycin-TCO, which has a PEG8-TCO attachment at the C-terminal carboxylic acid, retains the biological activity of the parent drug by evaluating its potency against *S. aureus*. A minimum inhibitory concentration (MIC) assay was performed with both vancomycin and vancomycin-TCO (**Figure S10A**). Vancomycin had an MIC of 1 μg/mL, which increased for vancomycin-TCO to 16 μg/mL. This increase is consistent with the limited range of PEG modified vancomycin compounds reported in the literature^46, 47^ (albeit more pronounced, possibly due to the longer PEG length). This result indicates that the TCO-PEG8-modified vancomycin retains the antibiotic activity of the parent drug and suggested that it may be effective at inhibiting bacterial growth at the surface, where a high local concentration of drug is found.

For planktonic growth inhibition studies, a 96-well plate was coated using DOPA-Tet and grafted with vancomycin-TCO. Mid-log phase bacteria were added to the wells and incubated overnight, followed by addition of resazurin to quantify cell growth (**Figure S10B**). Surfaces that were coated/grafted using DOPA-Tet/vancomycin-TCO inhibited the growth of *S. aureus*. In contrast, surfaces which were not grafted led to no growth inhibition. This indicated that the antibiotic activity of vancomycin-TCO was retained upon surface immobilization, and that the DOPA-Tet coating by itself was not antimicrobial. This result was corroborated by CLSM studies wherein coated microscopy dishes were similarly incubated with bacteria overnight and examined with PI/SYTO™ 9 staining and (**Figure 5A** and **Figure S11**). In this study, the coated/vancomycin-grafted sample clearly showed a much larger population of dead cells, and far fewer live cells, then the other samples.

**Figure 5.**
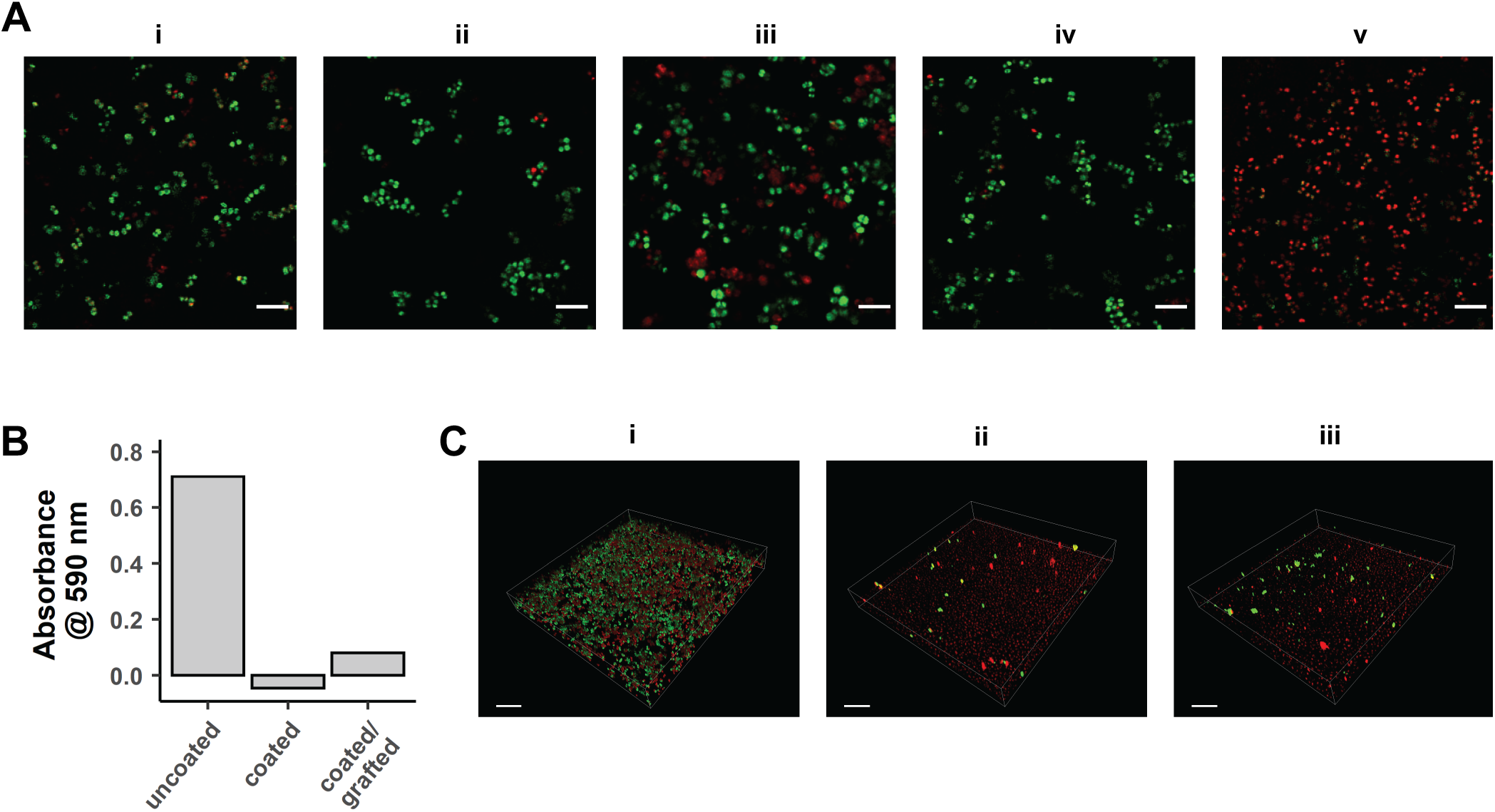
Imaging of planktonic and biofilm-forming *S. aureus* on functionalized surfaces. **A**. Confocal images of *S. aureus* cultures incubated on glass microscopy dishes that are incubated with (i) PBS, (ii) vancomycin, (iii) vancomycin-TCO, or coated with DOPA-Tet and then incubated with (iv) vancomycin or (v) vancomycin-TCO. Scale bar is 50 µm. **B**. Absorbance of uncoated, coated, or coated and vancomycin-grafted culture plates incubated for 72 hours with *S. aureus* and assayed with crystal violet. Shown as median values corrected for background absorbance with non-bacteria-containing samples. **C**. Confocal images of *S. aureus* biofilms grown on glass microscopy dishes that are (i) uncoated, (ii) coated, and (iii) coated/grafted with vancomycin-TCO. Scale bar is 20 µm. For both **A** and **C**, cultures were stained with a combination of PI and SYTO™ 9, which selectively label dead (red) and live (green) cells, respectively, and overlay images the channels are shown.

To further investigate the microbiological properties of the surfaces, we evaluated the impact of the surface functionalization on *S. aureus* biofilm formation. Toward this end, coated surfaces were incubated with *S. aureus* for 72 hours to allow for the formation of biofilms, then a solution of crystal violet (a common dye used as a quantitative indicator for biofilm formation)^48^ was introduced to the surfaces. After dissolution and measurement of absorbance at 590 nm, it was found that while a biofilm was formed on the uncoated surface, surfaces that were either coated using DOPA-Tet or coated/grafted using DOPA-Tet/vancomycin-TCO successfully inhibited biofilm formation (**Figure 5B** and **Figure S12**). To corroborate this result, coated microscopy dishes were similarly incubated with bacteria and examined with PI/SYTO™ 9 staining and CLSM. In this study, again no biofilm was observed on the coated or coated and vancomycin-grafted surfaces (**Figure 5C**).

## CONCLUSIONS

A central challenge in surface modifications directed at medical and biotechnology applications is to establish methods that chemoselective, generalizable, and fast. Bioorthogonal chemistries can open new possibilities in realizing such methods. In this work, we employed modern bioorthogonal approaches to enable bioinert and facile functionalization of material surfaces, leveraging the parallels of tyrosinase-catalyzed catecholamine polymerization and tetrazine ligation. Both reactions proceed rapidly and in the absence of a free transition metal. The method of tetrazine ligation for surface grafting was successful in functionalizing materials with an array of molecules possessing diverse function and structural complexity, including small molecules, enzymes, a peptide, and an antibiotic. The activity of these molecules was retained upon being grafted. Material properties were modified to produce surface-bound enzymatic catalysis, improved mammalian cell adhesion, and reduced bacterial adhesion and viability.

This work expands the scope of bioorthogonal chemistry^49^ by the implementation of previously underexplored chemoenzymatic approaches to surface chemistry. Through the development of molecular tools with superior properties, new and application-focused opportunities have thus become available to bioorthogonal reactions. The ease and flexibility with which materials and bioactive molecules can be integrated makes the described surface functionalization method well-positioned for enabling diverse biological applications.

## SUPPORTING INFORMATION

The Supporting Information includes chemicals, materials, and biologics; methods and characterization for syntheses; method details for experiments; and supplementary figures.

## ACKNOWLEDGMENTS

We thank Dr. Gene Hall for assistance with ATR-FTIR experiments and access to his instrument. We thank Dr. Spencer Knapp and Mark Dresel for allowing access to their polarimeter and Dr. Jacques Roberge for access to preparative HPLC. We thank Drs. Yan Cheng and Yufeng Zheng for their generous donation of the titanium substrates. We thank Drs. Zvi Loewy, Lisa Lyu, and Pragati Sharma for stimulating discussions. We thank Drs. Matthew Moschitto and Zheng Shi for helpful comments on the manuscript. This work was supported by the NIH / NIBIB Trailblazer Award (EB029548), ROI–HealthAdvance funding and NHLBI Award (U01HL150852), and New Jersey Health Foundation Research Grant Program (PC 88-20).

## AUTHOR CONTRIBUTIONS

K.H., Z.J., and E.C.I. conceived the study. K.H. and M.B. performed coating and grafting experiments. K.H., M.R.L.S., M.B., and T.A. established TCO attachment protocols. K.H. performed enzymatic assays. Z.J. conducted XPS experiments and mammalian cell studies. E.C.I., M.R.L.S, and M.B. performed the synthesis and characterization of new organic molecules. M.R.L.S. and M.B. conducted microbiological growth inhibition and biofilm assays. K.H. and E.C.I. wrote the manuscript with the help of Z.J., M.B., and M.R.L.S. E.C.I. supervised the research.

