## Supplementary Information for "Bioorthogonal Functionalization of Material Surfaces with Bioactive Molecules"

[§] These authors contributed equally

| Contents | Page |
|---|---|


| | |
|---|---|
| **Chemicals, Materials, and Biologics** | **S3–S4** |
| **General Synthetic Methods** | **S4** |
| **Preparation Procedures and Characterization Data for Small Molecules** | **S5–S14** |
| Compound ***S1*** | S5 |
| Compound ***S2*** | S6 |
| Compound ***DOPA-Tet*** | S7 |
| $^{1}H$ and $^{13}C$ NMR Spectra | S8–S13 |
| **Surface Functionalization** | **S14–S15** |
| **Preparation of TCO-containing MOIs** | **S16–S20** |
| $^{1}H$ and $^{13}C$ NMR Spectra of Vancomycin-TCO | S19–S20 |
| **Experimental Method Details** | **S21–S26** |
| The Activity of TCO-Enzymes vs. Native Enzymes | S21 |
| Grafting Verification via Small Molecule Fluorophore | S21 |
| Colorimetric Assay using ALP and Combination GOx / HRP | S22 |
| Stability of Grafted Enzymes After Repeated Use | S22 |
| Cell Culture | S23 |
| MTT Assay | S23 |
| Fibroblast Adhesion and Morphology on NanoECM | S23–S24 |
| Fibroblast ECM Deposition on NanoECM | S24 |
| Minimum Inhibitory Concentration Assay | S24 |
| Biofilm Crystal Violet Assay | S25 |
| Biofilm Microscopy Protocol | S25 |
| Planktonic Bacterial Growth Inhibition at the Surface | S26 |
| Live Cell Microscopy of Planktonic Bacteria on the Surface | S26 |
| **Surface Wettability and XPS** | **S27** |
| **ATR-FTIR Analysis** | **S28** |
| **Activity of Tyrosinase on *N*-Boc Protected DOPA-Tet** | **S29** |
| **Coating Stability at Various pH and in Human Serum** | **S30** |
| **Verification of Activity of TCO-Conjugated Enzymes** | **S31** |
| **Grafting Verification via Small Molecule Fluorophore** | **S32** |
| **Effect of Washing on ALP Activity** | **S33** |
| **Stability of Grafted ALP in Human Serum** | **S34** |
| **MTT Assay** | **S35** |
| **Microbiology** | **S36–S38** |
| Vancomycin Surface Inhibition of *S. aureus* Growth | S36 |
| CLSM images of *S. aureus* culture | S37 |
| Absorbance values for crystal violet assay | S38 |
| **References** | **S39** |

### Chemicals, Materials, and Biologics

**Reagents:** 3,4-Dihydroxy-L-phenylalanine (L-DOPA) was purchased from Alfa Aesar. 1-ethyl-3-[3-dimethylaminopropyl]carbodiimide HCl salt (EDC HCl), *N*,*N*-diisopropylethylamine (DIPEA), trifluoroacetic acid (TFA), di-tert-butyl dicarbonate ($Boc_2O$), *N*-methylmorpholine (NMM), para-nitro phenol phosphate, para-nitro phenol, trypsin, $N^{\alpha}$-Benzoyl-DL-arginine 4-nitroanilide hydrochloride, *N*-hydroxy succinimide (NHS), vancomycin (from *Streptomyces orientalis*), glutaraldehyde, paraformaldehyde, α-MEM medium, bovine serum albumin (BSA), agar, anti-vinculin monoclonal antibody, fluorescein isothiocyanate (FITC)-labeled goat anti-rabbit IgG, tetramethylrhodamine-isothiocyanate (TRITC)-phalloidin kit, 4′,6-diamidino-2-phenylindole dihydrochloride (DAPI), actin cytoskeleton/focal adhesion staining kit (FAK100), and Triton X-100 were purchased from Millipore-Sigma. 1-hydroxybenzotriazole (HOBt) was purchased from TCI America. (3-phenyl-6-Methyl-1,2,4,5-tetrazine)-PEG4-amine HCl salt (commercially named methyltetrazine-PEG4-amine HCl salt), TCO-PEG24-COOH, TCO-PEG8-$NH_2$, and TCO-PEG8-NHS were purchased from BroadPharm. Sulfo-Cy5-amine (Cy5) was purchased from Lumiprobe. Cy5-TCO was purchased from Click Chemistry Tools. Cyclo[Arg-Gly-Asp-D-Phe-Lys] (c(RGDfK)) was purchased from Apex Bio. 1-[Bis(dimethylamino)methylene]-1H-1,2,3-triazolo[4,5-b]pyridinium 3-oxide hexafluorophosphate (HATU) was purchased from CombiBlocks. 3-(4,5-dimethylthiazol-2-yl)-2,5-diphenyl tetrazolium bromide (MTT) cell proliferation assay kit was purchased from BioVision Inc. Dulbecco's Modified Eagle Medium (DMEM), Hanks' Balanced Salt Solution (HBSS), and Tryptic Soy Broth (TSB) were purchased from VWR International. Alkaline phosphatase (ALP), glucose oxidase (GOx), horseradish peroxidase (HRP), and tyrosinase were purchased from Millipore-Sigma. Propidium iodide (PI) was purchased from Dojindo, Japan. SYTO™ 9 and PrestoBlue™ were purchased from Thermo-Fisher Scientific.

**Buffers and Solvents:** Phosphate-buffered saline (PBS), 4-morpholineethanesulfonic acid (MES), tris(hydroxymethyl)aminomethane (Tris), Tris-buffered saline (TBS), and 2-[4-(2-hydroxyethyl)piperazin-1-yl]ethanesulfonic acid (HEPES) were purchased from Millipore-Sigma. Dry $CH_2Cl_2$ and dimethylformamide (DMF) were purchased from Millipore-Sigma. Dry methanol (MeOH), ethyl acetate (EtOAc), and diethyl ether ($Et_2O$) were purchased from Fisher Scientific. Deuterated solvents were purchased from either Cambridge Isotope Laboratories or Millipore-Sigma. Water was deionized and filtered to a resistivity of 18.2 ΩM with a Milli-Q® Plus water purification system (Millipore-Sigma). Buffers were prepared freshly in Milli-Q® water and their pH was adjusted using HCl or NaOH.

**Materials:** 700 nm fiber diameter NanoECM™ (a randomly oriented, electrospun polycaprolactone fiber product) was purchased from Nanofiber Solutions, LLC. Glass was purchased from VWR International. Si/$SiO_2$ wafer substrate was purchased from University Wafer. All of these materials were cleaned ultrasonically in ethanol and water for 15 min before use, except NanoECM, which was used as received. Commercially available pure titanium rods were cut into plates and polished up to 1200 grit using Silicon Carbide paper and then ultrasonically rinsed in acetone, ethanol, and water for 15 min each. Non-treated, pre-sterile 96-well tissue culture plates (VWR) were used for bacterial growth inhibition. glass bottom culture dish were purchased from Matsunami (D11130H).

**Biologics:** NIH3T3 cell line (CAL-1658) and *Staphylococcus aureus* (ATCC6538) were purchased from the American Type Culture Collection.

If not specified here, other chemicals, materials, and biologics will be detailed below.

### General Synthetic Methods

For the chemical synthesis of organic compounds, all reactions were performed under a dry nitrogen atmosphere unless otherwise stated. All glassware was oven-dried before use. Purification of the synthesized compounds was performed using a Büchi Reveleris® flash chromatography system equipped with either a silica gel or C18 column. Observed rotation ($\alpha_{obs}$) values were measured in a standard glass cell (100 mm, 1 mL) using a sodium D-line lamp at 20 ºC in a PerkinElmer Model 241 Polarimeter. Specific rotation [α] was calculated based on $[\alpha]^{20}_{D} = (\alpha_{obs})/[(g_{sample}$ in 1 mL) x 1 dm]. Nuclear Magnetic Resonance (NMR) spectroscopic analyses were carried out on a Bruker Avance Neo 500 MHz spectrometer. $^{1}$H NMR spectra were acquired at 500 MHz and $^{13}$C NMR spectra were acquired at 126 MHz. Chemical shifts in parts per million (δs) for $^{1}$H NMR spectra were referenced to $(C\boldsymbol{H}_3)_4Si$ at δ = 0.00 ppm, to $C\boldsymbol{H}D_2S(O)CD_3$ at δ = 2.50 ppm, to ***H***DO at δ = 4.79 ppm, or to $C\boldsymbol{H}Cl_3$ at δ = 7.26 ppm. $^{13}$C NMR spectra were referenced to $\boldsymbol{C}D_3S(O)\boldsymbol{C}D_3$ at δ = 39.52 ppm, to $\boldsymbol{C}D_3OD$ at δ = 49.00 ppm or to $\boldsymbol{C}DCl_3$ at δ = 77.16 ppm. The following abbreviations are used to describe NMR resonances: s (singlet), d (doublet), t (triplet), m (multiplet), dd (doublet of doublets), ddd (doublet of doublet of doublets), br (broad), and app (apparent). Coupling constants (*J*) are reported in Hz. Liquid chromatography followed by high-resolution mass spectroscopy (LC-HRMS) analysis in the ESI mode was carried out on a Waters Acquity-Xevo G2-XS QTof.

### Preparation Procedures and Characterization Data for Small Molecules

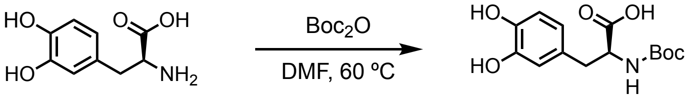


L-DOPA *S1*

**Synthesis of (*S*)-2-((tert-butoxycarbonyl)amino)-3-(3,4-dihydroxyphenyl)propanoic acid**, ***S1***: To a 50-mL round-bottom flask L-DOPA (0.5 g, 2.54 mmol, 1 equiv), $Boc_2O$ (1.11 g, 5.08 mmol, 2 equiv), and 9 mL of DMF was added. The heterogeneous mixture was purged with $N_2$ for 10 minutes and stirred at 60 °C for 24 hours under an $N_2$-enriched atmosphere. Upon cooling down to room temperature, the reaction mixture was added into a vigorously mixing emulsion of half-saturated brine (300 mL) and EtOAc (300 mL). The mixture was transferred into a separatory funnel after being stirred for 5 minutes. The organic layer was sequentially washed with half-saturated brine (300 mL) and brine (100 mL). The collected organic layer was dried over anhydrous sodium sulfate and concentrated under reduced pressure. The resulting oil was placed under high vacuum for 24 hours, which provided compound ***S1*** (0.7 g, 2.4 mmol, 92% isolated yield) as a solid with pearl-white color.

**$^1$H NMR** (500 MHz, $d_6$-DMSO) δ 8.72 (br s, 2H), 6.93 (d, *J* = 8.0 Hz, 1H), 6.61 (d, *J* = 2.0 Hz, 1H), 6.60 (d, *J* = 8.0 Hz, 1H), 6.47 (dd, *J* = 8.0, 2.0 Hz, 1H), 3.97 (ddd, *J* = 10.0, 8.0, 4.5 Hz, 1H), 2.80 (dd, *J* = 14.0, 4.5 Hz, 1H), 2.63 (dd, *J* = 14.0, 10.0 Hz, 1H), and 1.33 (s, 9H).

**$^{13}$C NMR** (126 MHz, $d_6$-DMSO) δ 173.8, 155.4, 144.9, 143.8, 128.7, 119.8, 116.5, 115.3, 78.0, 55.6, 35.9, and 28.2.

**HRMS** (ESI) m/z: Calculated for $[C_{14}H_{18}NO_6]^-$, $[M - H]^-$, requires 296.1140; found 296.1112.

**$[\alpha]^{20}_D$** = +11.8° (c = 0.03 g/mL, MeOH).

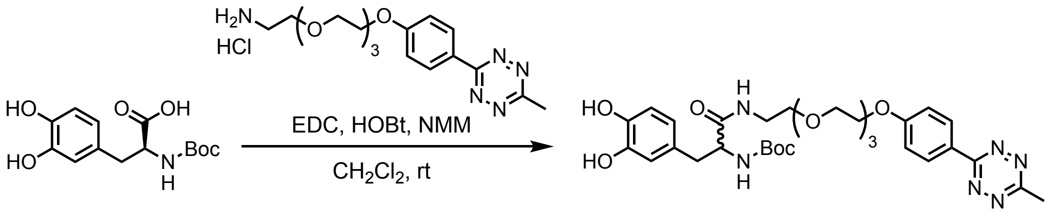


**Synthesis of tert-butyl (±)-(15-(3,4-dihydroxyphenyl)-1-(4-(6-methyl-1,2,4,5-tetrazin-3-yl)-phenoxy)-13-oxo-3,6,9-trioxa-12-azapentadecan-14-yl)carbamate, *S2*:** In a 20 mL amber vial, ***S1*** (150 mg, 0.50 mmol, 1 equiv), 3-phenyl-6-Methyl-1,2,4,5-tetrazine HCl salt (200 mg, 0.50 mmol, 1 equiv), EDC HCl (106 mg, 0.55 mmol, 1.1 equiv), HOBt (85 mg, 0.55 mmol, 1.1 equiv), and 5 mL of anhydrous $CH_2Cl_2$ were mixed and purged with $N_2$ for 10 minutes. Upon addition of NMM (61 µL, 0.55 mmol, 1.1 equiv) the via was sealed and the mixture was allowed to stir for 18 hours at room temperature. The reaction mixture was diluted in $CH_2Cl_2$ and extracted with a 5% aqueous citric acid solution. The aqueous phase was extracted with $CH_2Cl_2$ again and the combined organic layers were washed with brine, dried over sodium sulfate, and concentrated under reduced pressure. Silica gel column chromatography (MeOH/ $CH_2Cl_2$ step gradient) of the crude mixture gave ***S2*** (67% isolated yield) (215 mg, 0.33 mmol, 67% isolated yield) as a foam with magenta color.

**$^1$H NMR** (500 MHz, $CDCl_3$) δ 8.45 (d, *J* = 9.0 Hz, 2H), 7.01 (d, J = 9.0 Hz, 2H), 6.73 (d, *J* = 8.0 Hz, 1H), 6.71 (br s, 1H), 6.52 (d, *J* = 8.0 Hz 1H), 5.38 (br s, 1H), 4.25 (br s, 1H), 4.18 – 4.14 (m, 2H), 3.86 – 3.82 (m, 2H), 3.73 – 3.69 (m, 2H), 3.66 – 3.61 (m, 2H), 3.59 – 3.55 (m, 2H), 3.51 – 3.46 (m, 2H), 3.44 – 3.19 (m, 4H), 3.01 (s, 3H), 2.95 – 2.88 (app m, 1H), 2.78 (dd, *J* = 13.0, 8.0 Hz, 1H), and 1.37 (s, 9H).

**$^{13}$C NMR** (126 MHz, $CDCl_3$) δ 171.9, 166.7, 163.8, 162.4, 155.5, 144.4, 143.7, 129.8, 128.6, 124.4, 121.4, 116.6, 115.6, 115.3, 80.3, 70.9, 70.6, 70.5, 70.1, 69.65, 69.63, 67.6, 56.2, 39.4, 38.5, 28.4, and 21.1.

**HRMS** (ESI) m/z: Calculated for $[C_{31}H_{43}N_6O_9]^+$, $[M + H]^+$, requires 643.3086; found 643.3118.

**$[\alpha]^{20}_D$** ~ 0.

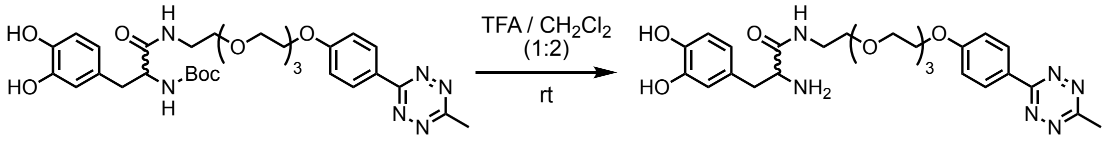


*S2* **DOPA-Tet**

**Synthesis of (±)-2-amino-3-(3,4-dihydroxyphenyl)-*N*-(2-(2-(2-(2-(4-(6-methyl-1,2,4,5-tetrazin-3-yl)phenoxy)ethoxy)ethoxy)ethoxy)ethyl)propanamide, *DOPA-Tet*:** In a 50 mL round-bottom flask, ***S2*** (215 mg, 0.34 mmol) was dissolved in 12 mL of $CH_2Cl_2$. After purging with $N_2$ for 10 minutes, 6 mL of previously $N_2$ purged TFA was added to the solution dropwise and the reaction mixture was allowed to stir at room temperature for 3 hours. The mixture was diluted first with $CH_2Cl_2$ and then chilled $Et_2O$, which resulted in precipitation of the product. The liquid was removed and the remaining precipitate was dried under reduced pressure, providing **DOPA-Tet** (150 mg, 0.23 mmol, 68% isolated yield) as a powder with magenta color. An aliquot was dissolved in water and lyophilized for the NMR spectral characterizations.

**$^1$H NMR** (500 MHz, $D_2O$) $\delta$ 8.15 (d, $J$ = 8.6 Hz, 2H), 7.06 (d, $J$ = 8.7 Hz, 2H), 6.69 (d, $J$ = 8.1 Hz, 1H), 6.54 (d, $J$ = 1.6 Hz, 1H), 6.49 (dd, $J$ = 8.1, 1.8 Hz, 1H), 4.26 – 4.21 (m, 2H), 4.03 (t, $J$ = 7.2 Hz, 1H), 3.94 – 3.89 (m, 2H), 3.80 – 3.75 (m, 2H), 3.74 – 3.70 (m, 2H), 3.68 – 3.65 (m, 2H), 3.61 – 3.56 (m, 2H), 3.54 – 3.50 (m, 1H), 3.41 (ddt, $J$ = 13.7, 9.8, 5.2 Hz, 2H), 3.24 (ddd, $J$ = 15.1, 7.2, 3.7 Hz, 1H), 2.98 (s, 3H), and 2.87 (app d, $J$ = 7.0, 2H).

**$^{13}$C NMR** (125 MHz, $CD_3OD$) $\delta$ 169.7, 168.1, 165.0, 164.0, 146.8, 146.1, 130.6, 126.8, 125.8, 121.8, 117.5, 116.7, 116.3, 71.7, 71.6, 71.5, 71.2, 70.7, 70.2, 68.9, 56.0, 40.5, 38.2, and 20.9.

**HRMS** (ESI) m/z: Calculated for $[C_{26}H_{35}N_6O_7]^+$, $[M + H]^+$, requires 543.2562; found 543.2599.

**$[\alpha]^{20}_D$** ~ 0.

***S1*** ($^1$H NMR: 500 MHz, $d_6$-DMSO)

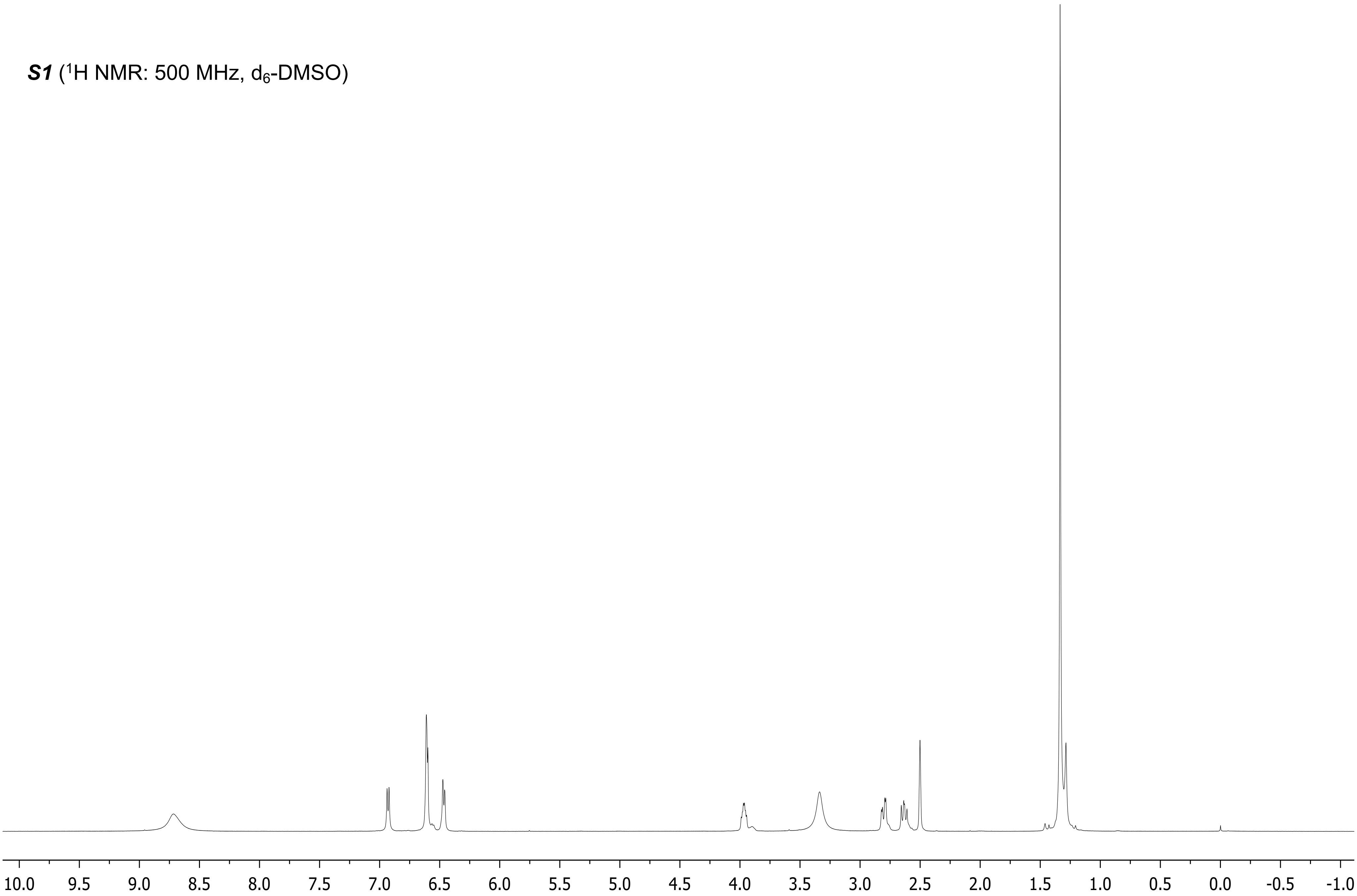

***S1*** ($^{13}C$ NMR: 126 MHz, $d_6$-DMSO)

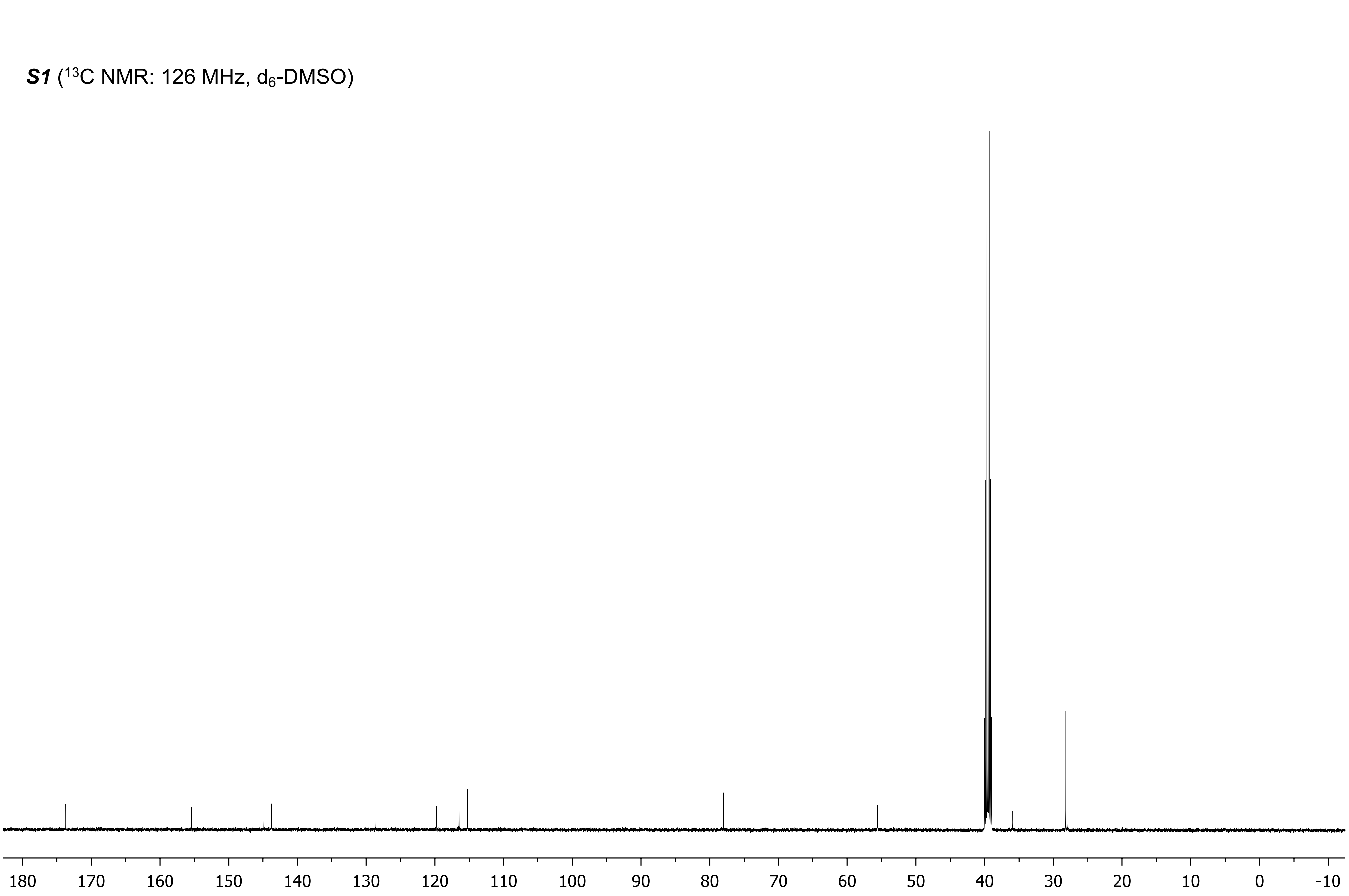

***S2*** ($^1$H NMR: 500 MHz, $CDCl_3$)

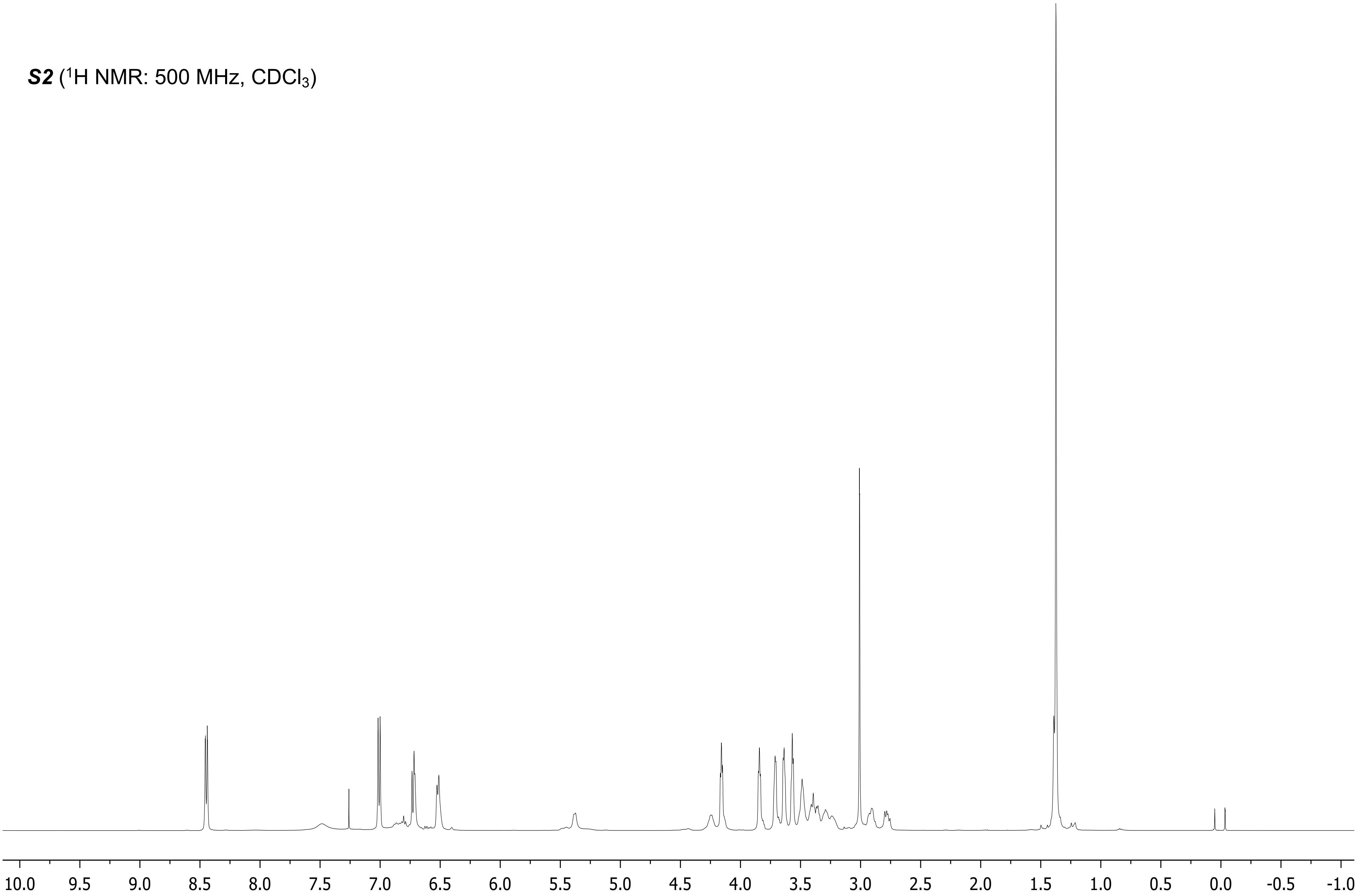

***S2*** ($^{13}$C NMR: 126 MHz, $CDCl_3$)

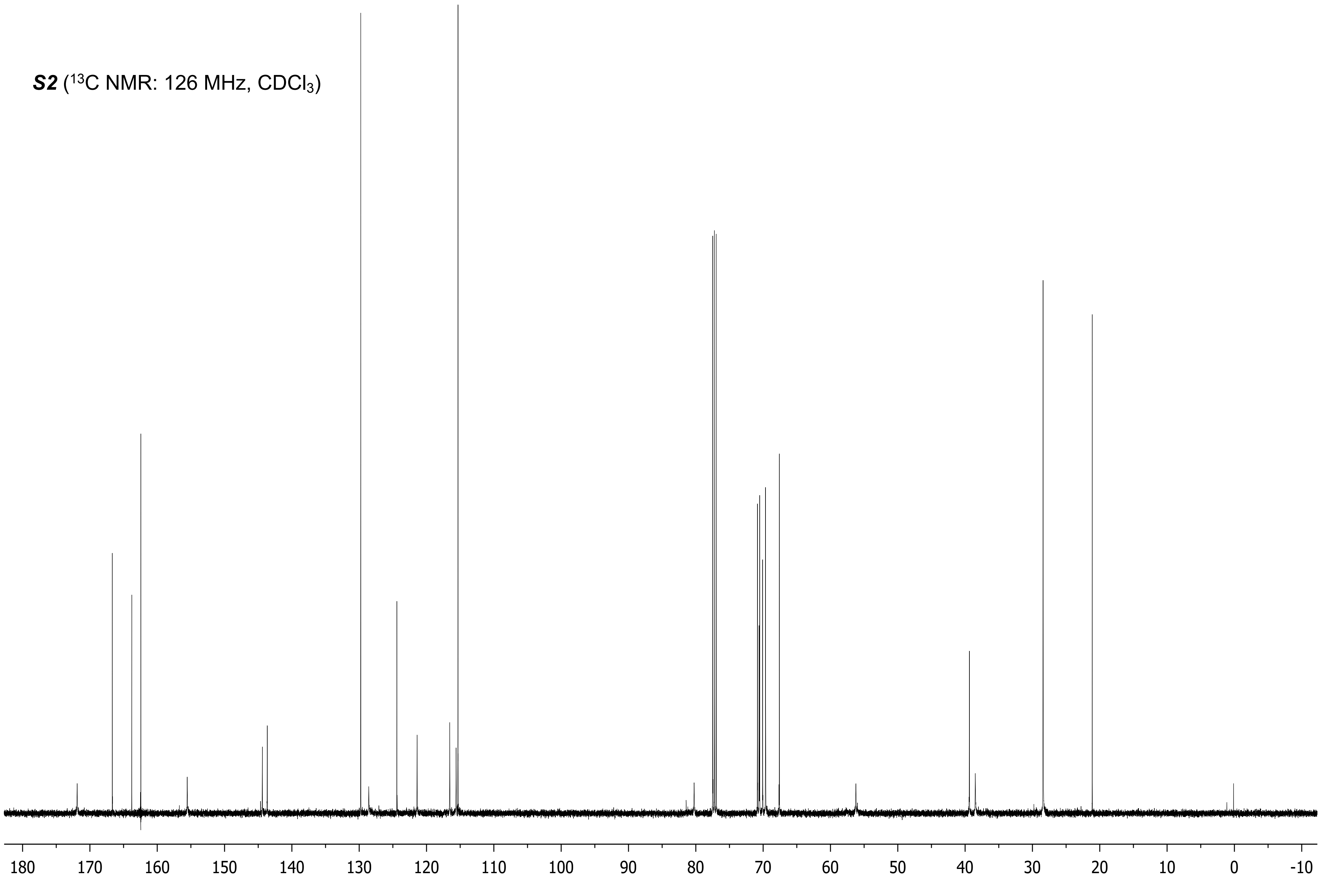

**DOPA-Tet** ($^1H$ NMR: 500 MHz, $D_2O$)

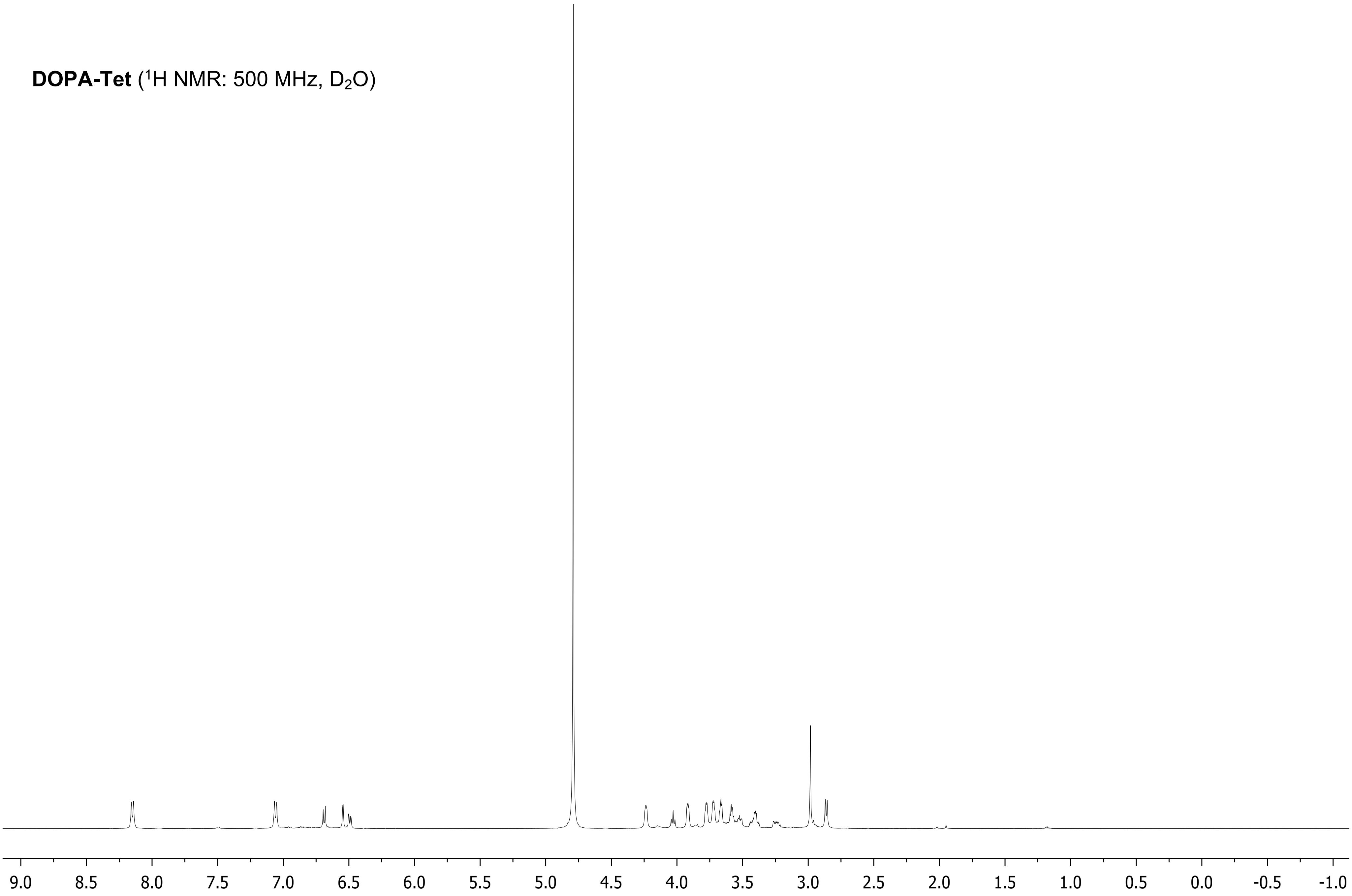

**DOPA-Tet** ($^{13}C$ NMR: 126 MHz, $CD_3OD$)

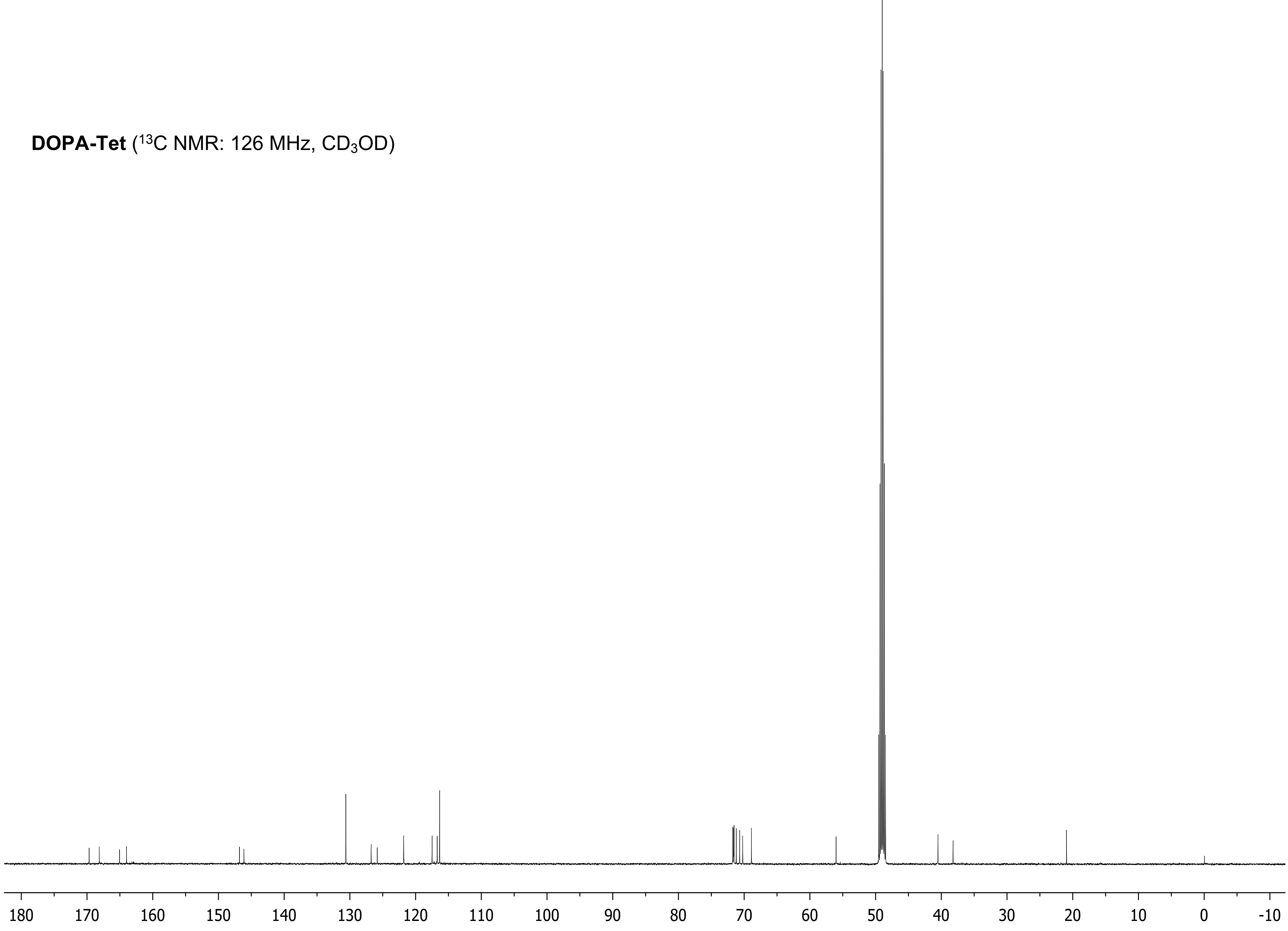

### Surface Functionalization

**General Methods:** Near-physiologic buffers were prepared for reactions involving biomolecules. PBS (pH 7.4) was prepared per the manufacturer's instructions. TBS was produced with 50 mM tris(hydroxymethyl)aminomethane and 150 mM NaCl and adjusted with HCl to pH 7.5 or 8.5.

For cell studies, buffers were sterilized by autoclaving, coating solutions were filtered using a 0.2 µm cellulose filter (VWR International), and coating procedures were performed in a CLASS II A2 Thermo Scientific biosafety cabinet.

Except where noted, all data analysis was done using R version 3.6.3 (packages used: tidyverse 1.3.0, broom 0.7.2, ggplot2 3.3.2). Welch's *t*-test was used for comparisons of sample means, as equal sample variances could not be assumed.

**Surface grafting method:** First, a solution of 1–10 mM DOPA-Tet and 2,500 U/mL tyrosinase in PBS was freshly prepared and immediately added to the surface. The solution was covered (to minimize evaporation) and incubated at room temperature for 1–2 hours, after which residual solution was rinsed off with PBS. After coating, a solution of 0.1–0.5 mM MOI-TCO in PBS was added to the coated surface, covered, and incubated for 1–2 hours at room temperature. After incubation, the residual solution was rinsed off with PBS. The coating and grafting solutions can be added either as a droplet or the surface can be submerged in the solution.

#### Characterization of Coating

*Tyrosinase Activity on DOPA-Tet:* As an initial test of the DOPA-Tet coating method, the plastic handle (polyethylene terephthalate) of a pH strip (Millipore-Sigma) and a polypropylene surface were coated with solutions of 10 mM L-DOPA or 10 mM DOPA-Tet in PBS, both with and without 2,500 U/mL tyrosinase. Samples were covered and incubated at room temperature for 1 hour before washing with DI water. Samples were imaged with a digital camera before and after rinsing.

*Surface Wettability:* Static contact angles on coated and uncoated titanium, silicon, and glass were measured using the sessile drop method. Samples of three materials, titanium discs prepared from a grade 2 (commercially pure) rod (McMaster-Carr), glass microscope slides (VWR International), and silicon wafers (University Wafer), were coated with 0.3, 1, 3, 10 mM DOPA-Tet or were left uncoated. Surfaces coated with above 10 mM were not tested due to potential confounding of the results by unevenness of the coatings. For each test, 5 µL aliquots of water were added at room temperature to the surface by slowly raising the surface to contact the suspended drop. 10-second videos were recorded on a benchtop goniometer (Ossila L2004A1) during contact and droplet spreading. Contact angles for each frame were calculated using Ossila Contact Angle v.3.1.1.0 and the mean of the last 5 frames, occurring at least 3 seconds after droplet settling, was used for static contact angle analysis. Three replicates of each coated or uncoated sample were prepared and measured.

*Coating characterization via X-ray photoelectron spectroscopy (XPS):* Three samples; a titanium disc, a glass microscope slide, and a silicon wafer as above; were coated with 10 mM DOPA-Tet. Three additional uncoated samples, one of each material, were also prepared. XPS studies were carried out on all six samples using a Thermo Scientific K-Alpha™ XPS System equipped with a monochromatic Al Kα source (h*v* = 1486.6 eV) at an energy step of 0.5 eV.

*Coating Characterization via ATR-FTIR:* Materials were characterized via ATR-FTIR to elucidate features of the structure. 10 µL of 10 mM DOPA-Tet, either with or without 2500 U/mL tyrosinase was deposited directly onto the ATR-FTIR diamond sensor. The tyrosinase-treated samples were washed with PBS to remove residual tyrosinase and unreacted DOPA-Tet. The samples without tyrosinase were not washed. All samples allowed to air dry. Additionally, a sample was produced where 10 µL of 10 mM (3-phenyl-6-methyl-1,2,4,5-tetrazine)-PEG4-amine was similarly deposited onto the sensor and allowed to air dry. Spectra of the samples were measured using a three-reflection diamond ATR attached to a PerkinElmer Spectrum 100 FT-IR. 32 scans were taken from 580 to 4000 $cm^{-1}$ at 4 $cm^{-1}$ resolution.

*Coating Stability at Various pH and in Human Serum:* Six 4 mL glass vials (VWR International) were coated with 10 mM DOPA-Tet and incubated for 5 days at 37 °C in 1 mL of either 50 mM MES pH 4.5 (Millipore-Sigma), 100 mM MES pH 6.0, PBS, TBS 8.5, 100 mM $NaHCO_3$ pH 9.5 (Millipore-Sigma) or 10% DMSO (Millipore-Sigma) in PBS. After incubation, samples were rinsed with water, dried, and imaged.

Two additional 4 mL glass vials were prepared and incubated in human serum. Both samples were coated with 10 mM DOPA-Tet, and one was grafted with 0.2 mM ALP-TCO. Both samples were incubated in 1 mL of human serum from clotted whole blood (Millipore-Sigma) for 5 days at 37 °C. After incubation, both samples were washed with PBS and 1 mL of 1 mg/mL p-NPP was added. After incubation for 1 hour at room temperature, both samples were imaged and their absorbance at 405 nm was measured on a spectrophotometer (Thermo Scientific NanoDrop One$^{C}$).

### Preparation of TCO-containing MOIs

**Preparation of ALP-TCO, GOx-TCO, and HRP-TCO:** Fresh aqueous solutions of EDC and NHS were prepared for this protocol, which was a prerequisite for its success. 20 µL of 200 mM TCO-PEG24-COOH was added to 400 µL of 100 mM EDC in 50 mM MES pH 6.0 (final concentrations of 9.5 mM TCO-PEG24-COOH and 95 mM EDC). The resulting mixture was incubated at room temperature (RT) for 15 minutes. After incubation, 400 µL of 200 mM NHS in 50 mM MES pH 6.0 was added to the solution, which was then incubated at room temperature for 30 minutes. To this solution 200 µL of either 0.344 mM ALP in 50 mM TBS pH 7.4, 1 mM GOx in 1x PBS pH 7.4, or 0.5 mM HRP in 1x PBS pH 7.4 was added. The solution was tumbled and incubated at room temperature for 2 hours to complete the reaction and produce the TCO-conjugated enzyme.

After reaction completion, 0.4 mL of the reaction solution was added to a 3 kDa cutoff centrifuge filter (UFC5003, Sigma) and centrifuged (Eppendorf Centrifuge 5430R) at 14,000 rcf for 15 minutes. The filtrate in the tube was discarded, and additional 0.4 mL aliquots of the reaction solution were added to the filter, centrifuged, and the filtrate discarded until all the reaction solution was filtered. Two successive aliquots of 0.4 mL water were added to the filter and centrifuged at 14,000 rcf for 15 minutes to wash away any remaining chemical reagents. The filtrate was discarded, the column was eluted with 0.5 mL of either 50 mM TBS (pH 7.5) for ALP or PBS (pH 7.4) for HRP or GOx, and the eluent was collected. An additional 0.5 mL of eluent was added and the eluted product fractions were combined, frozen at -80 °C overnight, and lyophilized (Labconco Freezone 4.5 Plus) at approximately 0.06 torr pressure. After lyophilization, either 50 mM TBS (pH 7.5) for ALP or PBS (pH 7.4) for HRP or GOx, was added to the tube to make a stock solution of the TCO-conjugated enzyme at the desired concentration (typically 0.5 mM). For calculations of the concentration of the final product, it was assumed that the enzyme was successfully conjugated and recovered quantitatively.

**Preparation of c(RGDfK)-TCO:**

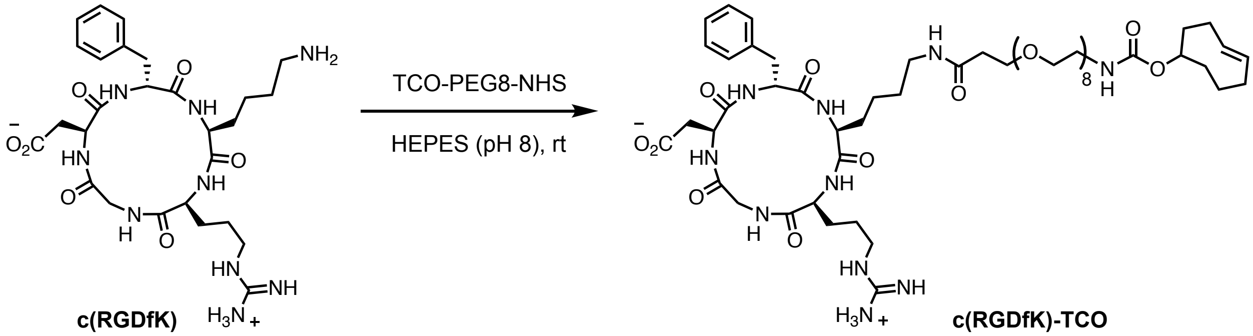


c(RGDfK) (1 mg, 1.66 μmol, 1 equiv) was combined with TCO-PEG8-NHS (2.5 mg, 3.62 μmol, 2.2 equiv) in 150 μL 50 mM HEPES (pH 8.0) and incubated at room temperature for 1 hour. The reaction product was confirmed by HR-LCMS and used in cell studies without any purification.

**HR-LCMS** (ESI) m/z: Calculated for $[C_{55}H_{91}N_{10}O_{18}]^+$, $[M + H]^+$, requires 1179.6507; found 1179.6511.

**Synthesis of Vancomycin-TCO:**

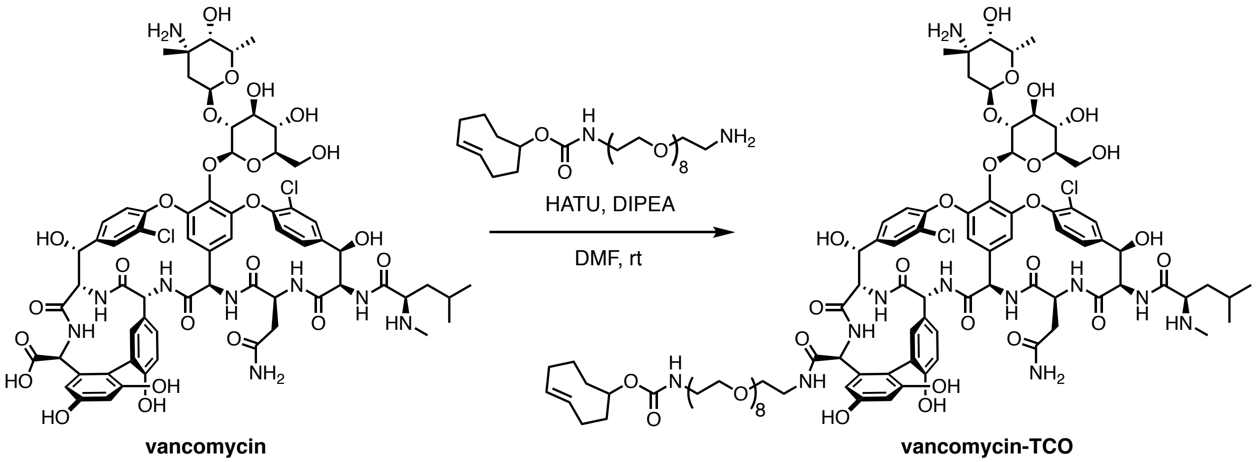


Vancomycin-HCl (90.8 mg, 0.061 mmol, 1 equiv) was suspended in 2 mL DMF under nitrogen, then DIPEA (31.9 μL, 0.183 mmol, 3 equiv) and HATU (69.7 mg, 0.183 mmol, 3 equiv) were added. The resulting yellow solution was stirred for 5 minutes, then TCO-PEG8-$NH_2$ (43.1 mg, 0.076 mmol, 1.3 equiv) dissolved in 1 mL DMF was added. The mixture was stirred overnight at room temperature, quenched with water, and directly concentrated under reduced pressure. The crude product was redissolved in water and purified using reverse-phase chromatography on C18 column (Eluent A: 0.1% TFA in $H_2O$, Eluent B: 0.1% TFA in ACN, step gradient from 100% A to

100% B). Product fractions (based on HRMS) were collected and lyophilized, providing vancomycin-TCO (46.1 mg, 38% isolated yield) as a pale-yellow solid. For NMR characterization and biofilm assays, this material was subjected to HPLC purification (same elution system as above). The NMR peak assignments were guided by a previously reported vancomycin characterization[1].

**$^1$H NMR** (500 MHz, $(CD_3)_2SO$) δ 9.31 (d, *J* = 3.2 Hz, 1H), 8.97 (br s, 1H), 7.86 (d, *J* = 1.0 Hz, 1H), 7.75 – 7.63 (m, 1H), 7.55 (d, *J* = 8.4 Hz, 1H), 7.50 (s, 1H), 7.46 (dd, *J* = 8.4, 1.3 Hz, 1H), 7.33 (d, *J* = 8.4 Hz, 1H), 7.28 (br s, 2H), 7.20 (s, 1H), 6.76 (d, *J* = 8.3, 1H) , 6.70 (dd, *J* = 8.3, 2.2 Hz, 1H), 6.65 (br s, 1H), 6.39 – 6.35 (m, 1H), 6.25 (app s, 1H), 5.97 (app s, 1H), 5.86 (d, *J* = 6.2 Hz, 1H), 5.76 (s, 1H), 5.62 – 5.53 (m, 2H), 5.43 (ddd, *J* = 14.6 Hz, *J* = 11.0 Hz, *J* = 3.5 Hz, 1H), 5.36 (d, *J* = 5.5 Hz, 1H), 5.28 – 5.21 (m, 2H), 5.18 (app s, 2H), 5.11 (br s, 1H), 4.94 (br s, 1H), 4.68 (q, *J* = 6.4 Hz, 1H), 4.44 (s, 1H), 4.39 (s, 1H), 4.28 – 4.17 (m, 2H), 4.06 – 4.00 (m, 1H), 3.94 (br s, 1H), 3.68 (d, *J* = 10.4 Hz, 1H), 3.59 – 3.46 (m, 28H)*, 3.39 – 3.35 (m, 2H), 3.27 (s, 2H), 3.17 (s, 1H), 3.13 – 3.03 (m, 2H), 2.92 (s, 1H), 2.64 (app br s, 2H), 2.34 – 2.08 (m, 4H), 1.95 – 1.80 (m, 4H), 1.30 (s, 3H), 1.69 – 1.60 (m, 3H), 1.60 – 1.49 (m, 3H), 1.49 – 1.41 (m, 1H), 1.30 (app s, 3H), 1.07 (d, *J* = 6.0 Hz, 3H), 0.91 (d, *J* = 6.0 Hz, 3H), 0.86 (d, *J* = 6.0 Hz, 3H).
*ethylene glycol units.

**$^{13}$C NMR** (125 MHz, $(CD_3)_2SO$) δ 171.44, 171.39, 171.37, 171.32, 170.32, 170.14, 169.02, 167.94, 158.01, 157.76, 157.52, 157.11, 156.27, 155.80*, 154.97, 152.57, 151.25, 148.16, 142.49, 139.65, 137.55, 135.55, 134.93*, 132.53*, 131.92, 129.65, 129.48, 127.29, 127.21, 126.30, 125.28, 124.36, 123.40, 121.18, 118.53, 116.14, 113.73, 107.35, 104.65, 101.98, 101.25, 96.75, 79.07*, 78.19, 76.99, 76.73, 74.42*, 69.77*‡, 69.72*, 69.63*, 69.52*, 69.16*, 69.14*, 68.92*, 63.10, 61.96, 61.88, 61.23, 61.12, 57.40, 54.84, 53.83, 53.78, 53.63, 50.94, 40.68, 38.19*, 33.75*, 33.70, 33.12, 32.16*, 31.26, 31.15, 30.59*, 29.00, 25.28, 25.14, 24.54, 23.69, 22.74, 22.51, 22.47, 22.43, 22.32, 21.93, 16.80.
* TCO-PEG8 group.
‡ 10 Cs (representing 5 repeating units of ethylene glycol) or more (with potential overlaps).

**HRMS** (ESI) m/z: Calculated for $[C_{93}H_{127}Cl_2N_{11}O_{33}]^{2+}$, $[M + 2H]^{2+}$, requires 997.8987; found 997.9009.

**vanco-TCO** ($^{1}H$ NMR: 500 MHz, $d_6$-DMSO)

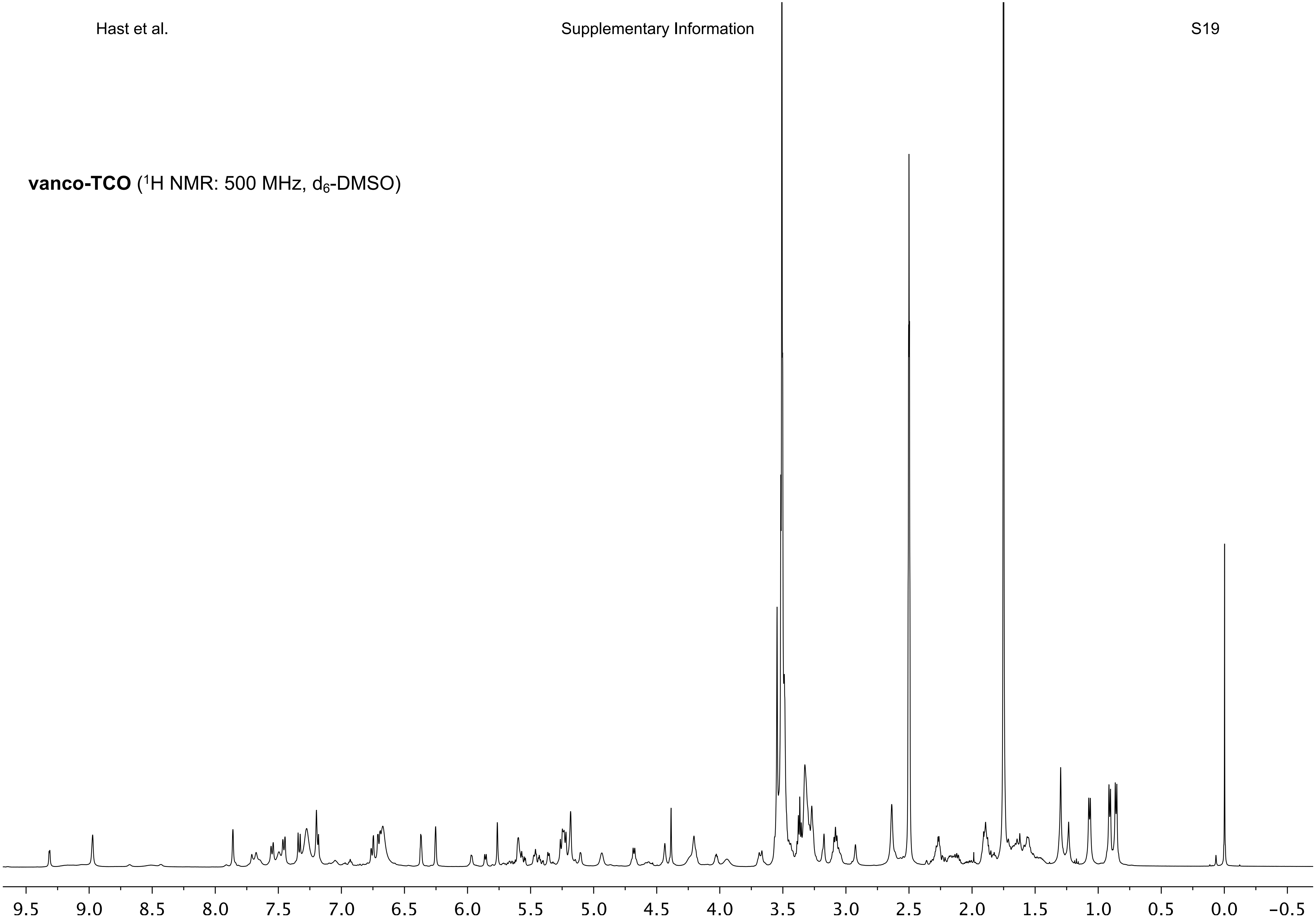

**vanco-TCO** ($^{13}C$ NMR: 126 MHz, $d_6$-DMSO)

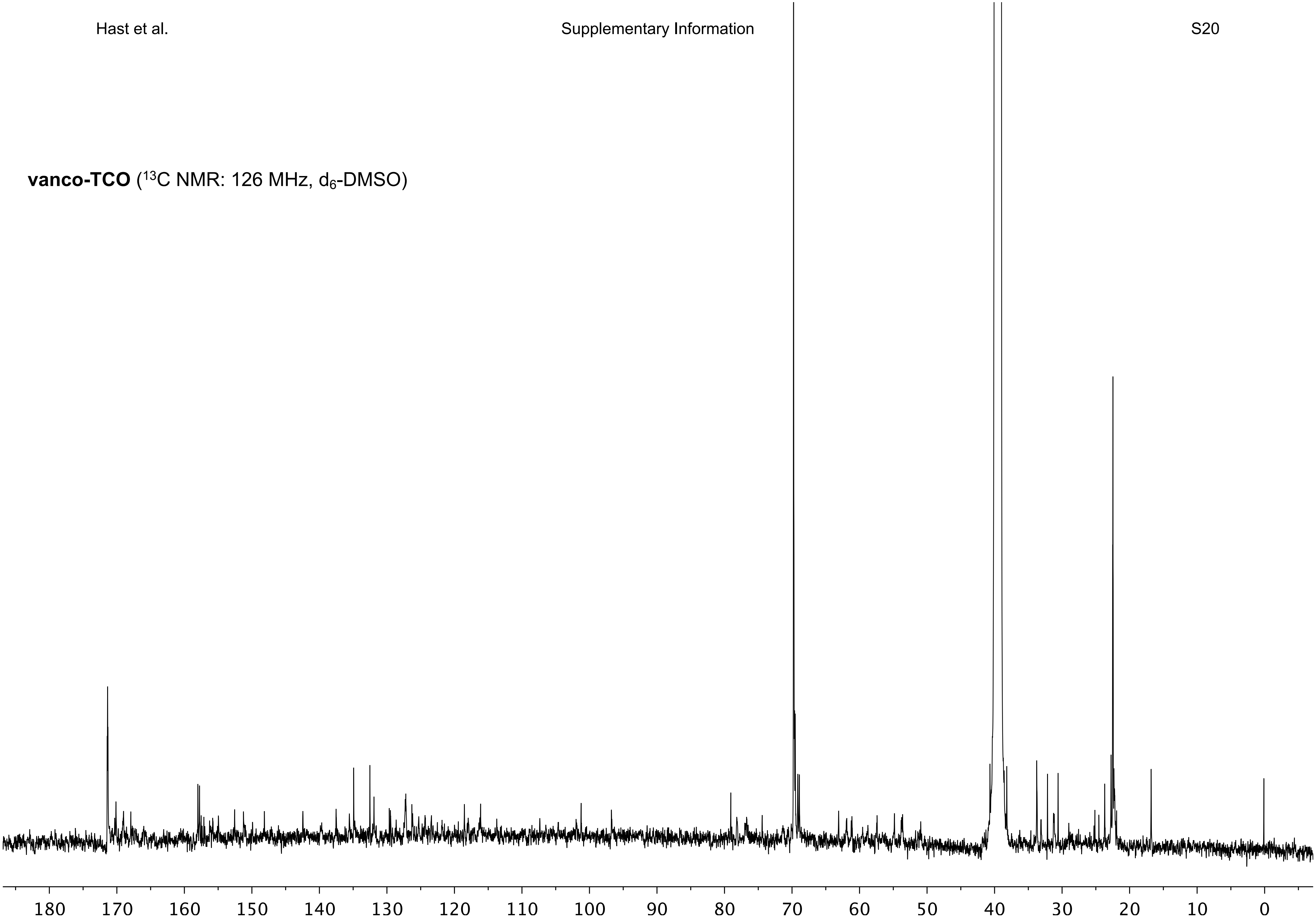

### Experimental Method Details

*The Activity of TCO-Enzymes vs. Native Enzymes:* To verify that TCO conjugation did not disrupt enzyme activity, we first prepared three solutions of **1)** 5 nM ALP-TCO and 1 mg/mL p-NPP (Millipore-Sigma), **2)** 5 nM ALP-TCO, and **3)** 1 mg/mL p-NPP, all in TBS 8.5. We similarly prepared a solution of 5 nM GOx-TCO, 5 nM HRP-TCO, 1 mM D-glucose (Research Products International), and 1 mM ABTS (Millipore-Sigma) in PBS, and four solutions where one of these four components was missing. All solutions were incubated at room temperature for 1 hour and their color changes were assessed visually.

We further compared the kinetics of the native and TCO-conjugated enzymes quantitatively using two UV-Vis assays. We first measured the kinetics of 100 µL 5 nM ALP, either TCO-conjugated or native, with 2.282–0.080 mM p-NPP in TBS 8.5 (16 total trials produced with 1.25x serial dilution of the substrate). We then measured the kinetics of 5 nM GOx and 5 nM HRP, both TCO-conjugated or both native. A 1.25x serial dilution was performed to produce 21 total trials of 100 µL, with concentrations of 32–0.369 mM ABTS and the same concentrations of D-glucose in PBS. Both assays produced an increase in absorbance at 405 nm, the evolution of which at room temperature was measured on a microplate reader (Molecular Devices SpectraMax iD3) over 1 hour. Small irregularities in the curves of some of the trials were observed in the first 2.5 minutes, so enzyme reaction rates were calculated using data from minutes 2.5–12, which included 19 data points per assay. Velocities were estimated by linear regression, and Michaelis-Menten parameters were estimated using non-linear least squares regression. R-squared values for all regression were above 0.95.

*Grafting Verification via Small Molecule Fluorophore:* Titanium discs were grafted with Cy5 or Cy5-TCO, both with and without first coating with DOPA-Tet (**Figure S6**). First, titanium discs were either coated with 10 mM DOPA-Tet or left uncoated. After rinsing, the discs were incubated with either 0.2 mM Cy5 or Cy5-TCO. For imaging, the slides were covered with a glass cover slip. The samples were imaged on a BioRad Chemidoc MP gel imager before and after rinsing off the fluorophore solution. For the rinsed samples 5 µL of PBS was added to the dried, coated disc to promote fluorescent activity. Water and PBS were found to be ineffective in removing residual Cy5 from the surface, and the samples were rinsed additionally with methanol.

All samples had similar fluorescence before rinsing, however, the sample that was coated and grafted with Cy5-TCO was detected to exhibit significantly higher fluorescence intensity than either the uncoated samples or the Cy5 incubated samples. The coated sample that was functionalized with Cy5 showed a small amount of fluorescence, suggesting non-specific binding between the fluorophore and the coating.

*Colorimetric Assay using ALP and Combination GOx / HRP:* To demonstrate the effectiveness of grafting complex biomolecules, we prepared assays using grafted enzymes from two different colorimetric assays, specifically ALP and a combination of GOx and HRP.

For grafting ALP, TBS was chosen as the buffer for washing and testing, because the high concentration of phosphate in PBS could affect the phosphatase activity of ALP. ALP-TCO was freshly prepared as described above. 0.2 mL microcentrifuge tubes were coated with 50 μL of 10 mM DOPA-Tet. Samples produced for visual comparison only (see **Figure 3C**) were coated with 100 μL. The tubes were washed thoroughly with TBS (pH 7.5) and sonicated to remove any loose aggregates. The tubes were grafted with 50 μL of either 0.2 mM ALP-TCO or 0.2 mM ALP, both in TBS. The tubes were washed thoroughly with TBS (pH 7.5) before 50 μL of 1 mg/mL p-NPP in TBS (pH 8.5) was added to each tube and incubated at room temperature for 1 hour. Absorbance at 405 nm was measured on a spectrophotometer (Thermo Scientific NanoDrop One$^{C}$). Three replicates for each grafted molecule, ALP-TCO or ALP, were prepared and measured.

For grafting GOx / HRP, both GOx-TCO and HRP-TCO were freshly prepared as described above. 0.2 mL microcentrifuge tubes were coated with 50 μL of 10 mM DOPA-Tet. Samples produced for visual comparison (see **Figure 3C**) were coated with 100 μL. The tubes were washed with PBS and sonicated to help remove any loose aggregates. The tubes were grafted with 50 μL of either 0.2 mM GOx-TCO and 0.2 mM HRP-TCO, or 0.2 mM GOx and 0.2 mM HRP. 50 μL of 1 mM D-glucose and 1 mM ABTS was added to each tube and were incubated at room temperature for 1 hour. Absorbance at 405 nm was measured on a spectrophotometer (Thermo Scientific NanoDrop One$^{C}$). Three replicates of each set of enzymes, TCO-conjugated or native, were prepared and measured.

*Stability of Grafted Enzymes After Repeated Use*: Eight 0.2 mL microcentrifuge tubes were coated with 50 μL of 10 mM DOPA-Tet. The tubes were washed thoroughly with TBS (pH 7.5). Three tubes each were grafted with 50 μL of either 0.1 mM ALP-TCO or 0.1 mM GOx-TCO and 0.1 mM HRP-TCO. The two remaining coated tubes were not grafted with any enzymes. The tubes were washed thoroughly with TBS (pH 7.5). 25 μL of 1 mg/mL p-NPP in TBS (pH 8.5) was added to the ALP-grafted tubes and 25 μL 1 mM ABTS and 1 mM D-glucose in PBS (pH 7.4) was added to the GOX/HRP grafted tubes. Each substrate solution was also added to one of the tubes that was not grafted (for controls). All eight tubes were incubated at room temperature for 1 hour. Absorbance at 405 nm was measured on a spectrophotometer (Thermo Scientific NanoDrop One$^{C}$). All tubes were then thoroughly washed with TBS (pH 7.5). Then TBS (pH 8.5) was added to the ALP-grafted tubes and PBS (pH 7.4) was added to the GOX/HRP grafted tubes. Each buffer solution was also added to one of the tubes that was not grafted. All eight tubes were incubated at room temperature for 1 hour after which absorbance at 405 nm was again measured. The buffer solutions were then removed from the tubes. This process of incubating with substrate and then buffer solutions was repeated for all tubes 9 more times, for a total of 10 cycles.

*Cell Culture:* NIH3T3 cells (CAL-1658, ATCC) were cultured in DMEM (VWR International) that contained 10% (v/v) fetal bovine serum (FBS) (Hyclone Laboratories, Inc.), 100 units/mL penicillin (VWR International), and 100 µg/mL streptomycin (VWR International). Cultivation was performed in a humidified VWR Air Jacketed Incubator with 5% $CO_2$ at 37 °C. The medium was refreshed every 2–3 days. For cell seeding, sub-confluent cells were harvested using 0.05% trypsin-EDTA, centrifuged, and resuspended to the desired density.

*MTT Assay:* Four samples of NanoECM were coated with 10 mM DOPA-Tet and ligated with 0.5 mM c(RGDfK)-TCO in PBS as described above. Four additional uncoated samples were prepared. The uncoated and coated samples were each incubated in serum-free DMEM at 37 °C for 72 hours with an extraction ratio of 1.25 cm$^2$/mL. The collected extracts were preserved at 4 °C and supplemented with 10% (v/v) FBS before use. NIH3T3 cells ($5\times10^4$ cells/mL) were seeded in 96-well tissue culture polystyrene plates (TCPS) and cultured for 24 hours to allow complete attachment. After seeding, the medium was replaced with an equal volume of extract and cultured for 1 day or 3 days. The normal DMEM and 10% DMSO-containing DMEM were set as negative and positive controls, respectively. After culturing, a colorimetric MTT assay (Biovision Incorporated) was performed according to the manufacturer's instructions. Briefly, the culture medium was discarded, and to each well 50 µL of serum-free α-MEM and 50 µL of MTT solution were added. The plates were incubated at 37 °C for 3 hours to yield formazan crystals. The formazan was dissolved in an MTT solvent under gentle shaking (Corning LSE Benchtop Shaking Incubator) in darkness, and its absorbance was measured at 590 nm using the microplate reader.

*Fibroblast Adhesion and Morphology on NanoECM:* For sterility, coating solutions were filtered using a 0.2 µm cellulose filter, and coating procedures were performed in a standard biosafety cabinet (Thermo Fisher Scientific, USA). Samples of 700 nm fiber diameter NanoECM (Nanofiber Solutions, LLC) discs were coated with either 3 mM or 10 mM DOPA-Tet and clicked with a solution of 0.5 mM c(RGDfK)-TCO in PBS. NIH3T3 cells ($5 \times 10^4$ cells/mL) were seeded onto uncoated and coated NanoECM substrates in 24-well TCPS plates and cultured at 37 °C for 6 hours. After culturing, the cells were fixed in 4% paraformaldehyde, permeabilized in 0.1% Triton X-100, and stained with a FAK100 kit (Millipore-Sigma) per manufacturer instructions. In brief, the cultures were blocked with 1% BSA in PBS for 30 minutes and incubated with an anti-vinculin monoclonal antibody (1:500 dilution) at room temperature for 1 hour. Subsequently, they were stained with FITC-conjugated goat anti-mouse IgG (1:100 dilution; 1 hour), TRITC-conjugated phalloidin (1:500 dilution; 1 hour), and DAPI (1:1000 dilution; 5 minutes). The samples were thoroughly washed with PBS and imaged using confocal laser scanning microscopy (CLSM, Zeiss LSM 710). Images were processed using either ZENBlack or ImageJ. The F-actin cytoskeleton (via TRITC-phalloidin), vinculin (via anti-vinculin), and nuclei (via DAPI) were visualized as red, green, and blue, respectively.

*Fibroblast ECM Deposition on NanoECM:* Five samples of NanoECM were coated with 10 mM DOPA-Tet and clicked with 0.5 mM c(RGDfK)-TCO in PBS as described above. Five additional uncoated samples were prepared. NIH3T3 cells ($5 \times 10^4$ cells/mL) were seeded onto each sample and incubated at 37 °C for up to 3 days. At 1, 4, 6, and hours and 1 and 3 days, a sample was gently rinsed with PBS, fixed overnight using 2.5% glutaraldehyde solution at 4 °C, and subsequently dehydrated in an ethanol gradient (50–100%) for 15 minutes. The resultant samples were dried in open air in the biosafety cabinet and sputter-coated with gold before imaging. Samples were imaged via scanning electron microscopy (Zeiss Sigma Field Emission SEM) at an accelerating voltage of 2 keV under vacuum.

*Minimum Inhibitory Concentration Assay:* A glycerol stock of *S. aureus* was struck on a tryptic soy agar plate and incubated overnight at 37 °C. Single colonies were then picked and diluted into tryptic soy broth (TSB) and left to grow overnight. In the morning, the culture was diluted approximately 40-fold into fresh TSB and grown to mid-log phase ($OD_{600}$ = 0.4 to 0.8). 5 mg/mL stock solutions of Vancomycin-TCO and vancomycin in DMSO were serially diluted two-fold with TBS across the wells of a non-treated 96-well plate, with concentrations ranging from 0.0625 to 128 µg/mL. The mid-log phase cultures were diluted to a final concentration of $5 \times 10^5$ colony forming units (CFU)/mL in TSB then 50 µL was added to each antimicrobial-containing well, giving a final antimicrobial concentration range of 0.031 to 64 µg/mL. All the plates were covered and incubated at 37 °C for 18 to 24 h. The MIC was defined as the lowest compound concentration at which no bacterial growth was visible (n = 3).

*Biofilm Crystal Violet Assay:* 100 µL of freshly prepared coating solution containing 10 mM DOPA-Tet and 2500 U/mL tyrosinase in PBS was added to the wells of a 96-well plate and incubated at room temperature for 6 hours. The wells were then washed three times with PBS. 100 µL of 0.2 mM vancomycin-TCO in PBS was added and the plate was incubated at room temperature for 1 hour and subsequently washed with PBS.

As above, *S. aureus* was grown overnight in TSB, then diluted 1:100 in TSB containing 3% glucose. 300 µL of the dilute bacteria was added to the coated well plate, which was then incubated at 37 °C for 72 hours. After incubation the media was decanted, and the plate was air dried for 5 minutes. 200 µL of 0.9% NaCl in sterile water was gently added to the wells, then decanted, and air dried for 5 minutes. The wells were washed twice more with 0.9% NaCl, and after the final drying, 200 µL methanol was added to each well. The plate was left at room temperature for 15 minutes, then the methanol was decanted and the plate air dried for 5 minutes. 100 µL of 3% crystal violet in sterile water was added to the wells and incubated at room temperature for 20 minutes. The dye was decanted, and the plate air dried. The wells were washed three times with 0.9% NaCl as above. After the final air drying, 150 µL of methanol was added to the wells, and mixed to allow for dissolution of the crystal violet. The dye solution was then decanted and diluted 1:10 in methanol, then the absorbance at 590 nm was recorded.

*Biofilm Microscopy Protocol:* 200 µL of freshly prepared coating solution containing 10 mM DOPA-Tet and 2500 U/mL tyrosinase in PBS was added to a glass bottom culture dush. The dish was incubated at room temperature for 6 hours, during which a coating was formed. The resulting coated surface was washed three times with PBS and 200 µL of 0.2 mM vancomycin-TCO in PBS was added. Then the surface was incubated at room temperature for 1 hour and subsequently washed with PBS.

*S. aureus* was grown overnight in TSB, then diluted 1:100 in TSB containing 3% glucose. 300 µL of the dilute bacteria was added to the microscope dishes, which were then placed in a 37 °C incubator. The samples were incubated for 72 hours, with the media being topped up every day to account for evaporation. The media was decanted and then the dishes were washed with 200 µL of 0.9% NaCl, followed by a 1-hour incubation at room temperature with 250 µL of TSB containing Syto9 (10 µM) and PI (10 µM). The dyes were then decanted and the samples were washed again with saline. Finally, 150 µL of HBSS was added, and the dishes were imaged on a Leica SP8 confocal microscope.

*Planktonic Bacterial Growth Inhibition at the Surface:* The wells of a non-treated clear 96 well culture plate were either left uncoated or coated with 10 mM DOPA-Tet and then incubated with either vancomycin-TCO, vancomycin, or PBS. A stock of *S. aureus* was grown to mid-log phase ($OD_{600}$ = 0.4 to 0.8) as above, then diluted to 5 x $10^5$ CFU/mL in TSB. 100 µL of culture was added to the wells and the plate was covered and incubated at 37 °C for 18 hours, after which 10 µL of PrestoBlue™ was added to each well. After 1 hour incubation, the media was transferred to clean wells and absorbance at 570 nm was recorded using a plate reader.

*Live Cell Microscopy of Planktonic Bacteria on the Surface:* Sterile glass-bottomed 35 mm dishes were either left uncoated or with 10 mM DOPA-Tet and then incubated with PBS or vancomycin-TCO. A stock of *S. aureus* was grown to mid-log phase ($OD_{600}$ = 0.4 – 0.8) as above, then diluted to ~5x$10^5$ CFU/mL in TSB and 250 µL was added to each dish. The plate was incubated at 37 °C for 18 hours, after which the media was decanted. A solution of propidium iodide (PI, 20 µM) and SYTO 9 (3.34 µM) was prepared in sterile Hanks' Buffered Salt Solution (HBSS) and 100 µL was added to each dish. The dishes were incubated for 30 mins in the dark at room temperature, after which the staining solution was decanted and 100 µL of HBSS was added to the dishes. Each dish was then imaged using a 63x oil objective on a Leica TCS SP8 confocal microscope.

### Surface Wettability and XPS

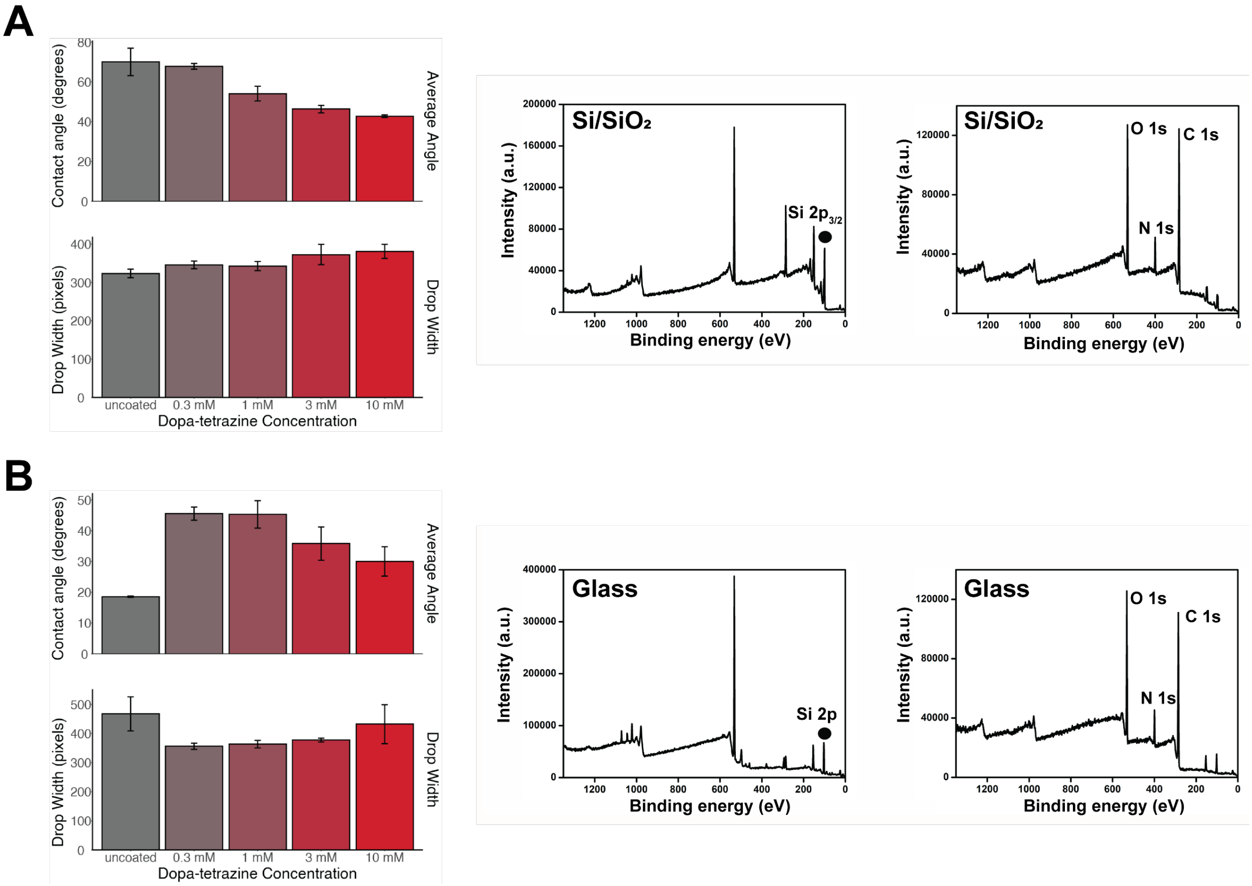


**Figure S1**. Coating characterization for uncoated and DOPA-Tet coated surfaces. (Left) Sessile Drop and (Right) XPS results for **(A)** silicon and **(B)** glass surfaces.

### ATR-FTIR Analysis

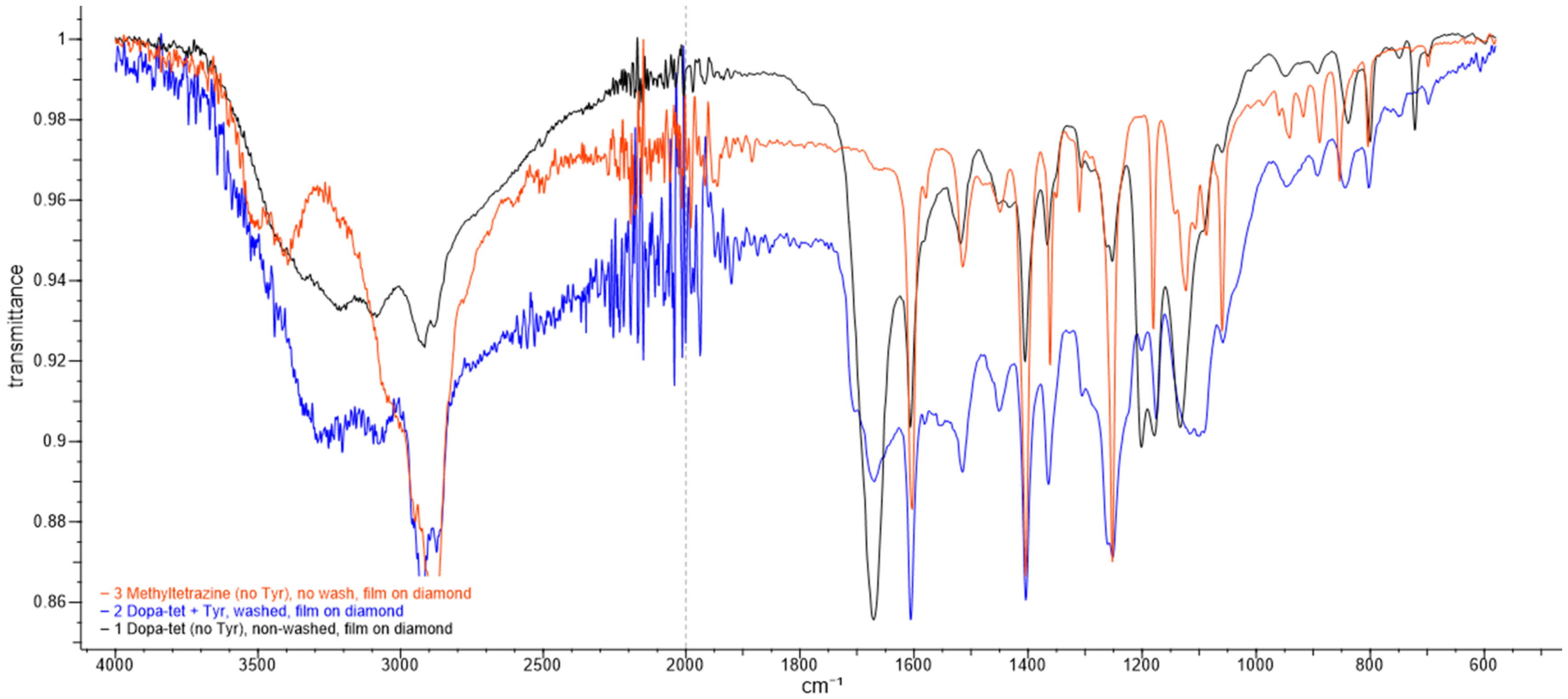


**Figure S2**. Attenuated total reflectance Fourier-transform infrared spectroscopy (ATR-FTIR) Spectra for coating materials. Spectra are shown for (Black) Dopa-Tet, (Blue) DOPA-Tet treated with tyrosinase, and (Orange) (3-phenyl-6-Methyl-1,2,4,5-tetrazine)-PEG4-amine (referred to in the manuscript as Tet-PEG4-NH2). Spectra were collected from solutions deposited directly onto the ATR-FTIR diamond sensor.

### Activity of Tyrosinase on *N*-Boc Protected DOPA-Tet (Compound *S2*)

Two samples of 10 mM compound ***S2*** were prepared in microcentrifuge tubes. 2,500 U/mL tyrosinase was added to one of the samples. A third sample of 10 mM DOPA-Tet with 2,500 U/mL tyrosinase was produced for visual comparison. The three samples were incubated at room temperature for 16 hours, and observations were made starting 2 hours after incubation.

No change in color, formation of aggregates, or coating of the tube was observed in the ***S2*** sample with tyrosinase, which remained indistinguishable from the ***S2*** sample without tyrosinase (**Figure S3**). As was typical, color change, aggregation, and deposition of material on the tube were observed in the DOPA-Tet sample with tyrosinase.

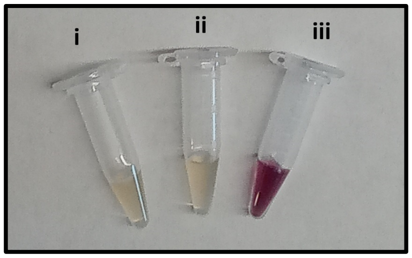


**Figure S3**. Activity of tyrosinase on compound ***S2*** and DOPA-Tet after 16 hours. **(i)** 10 mM ***S2***, **(ii)** 10 mM ***S2*** with tyrosinase, **(iii)** 10 mM DOPA-Tet with tyrosinase.

### Coating Stability at Various pH and in Human Serum

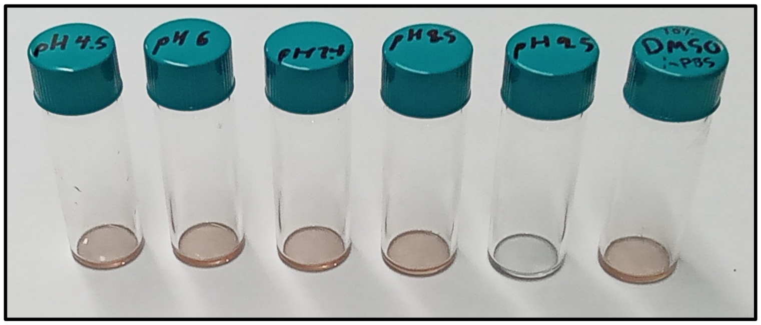


**Figure S4**. DOPA-Tet coatings after incubation for 5 days at 37 °C in various buffers. Buffers from left to right: 50 mM MES pH 4.5, 50 mM MES pH 6.0, 1x PBS pH 7.4, 50 mM TBS pH 8.5, 100 mM $NaHCO_3$ pH 9.5, 10% DMSO in 1x PBS pH 7.4.

### Verification of Activity of TCO-Conjugated Enzymes

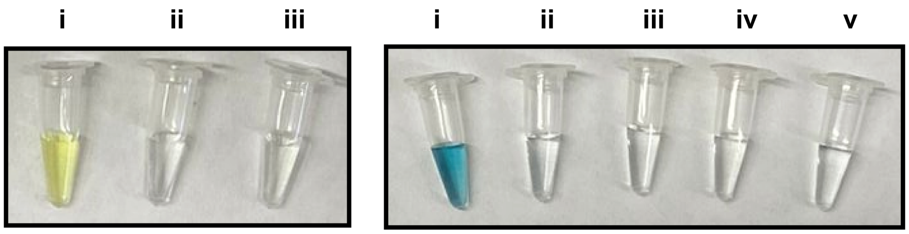


**Figure S5.** Colorimetric assays verifying components for coated assays. (Left) (i) ALP-TCO, p-NPP; (ii) ALP-TCO; (iii) p-NPP; (Right) (i) GOx-TCO, HRP-TCO, D-glucose, ABTS; (ii) same as (i) less GOx-TCO; (iii) same as (i) less HRP-TCO; (iv) same as (i) less D-glucose; (v) same as (i) less ABTS.

### Grafting Verification via Small Molecule Fluorophore

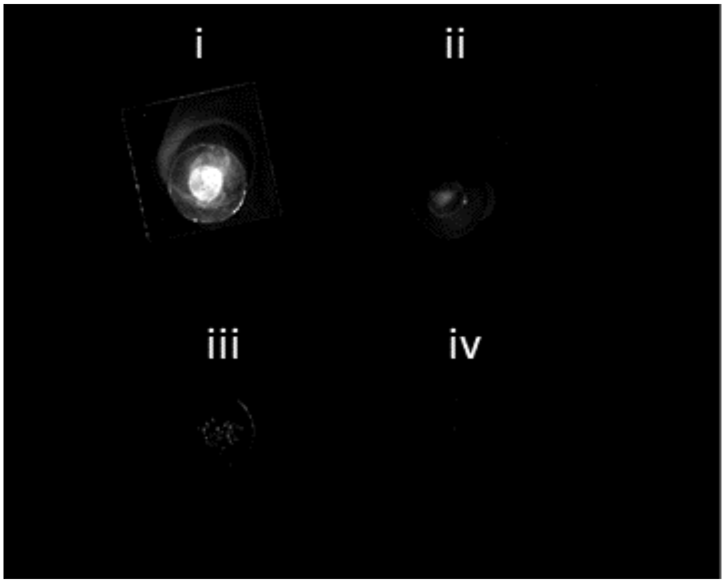


**Figure S6.** Cy5 and Cy5-TCO incubated on uncoated and DOPA-Tet coated titanium discs. **(i)** Cy5-TCO on DOPA-Tet coating. **(ii)** Cy5 on DOPA-Tet coating. **(iii)** Cy5-TCO on uncoated titanium. **(iv)** Cy5 on uncoated titanium.

### Effect of Washing on ALP Activity

A test was performed to examine the efficacy of various solvents and buffers in removing native ALP from DOPA-Tet coated surfaces. Seven wash solutions were prepared: **1)** Water, **2)** 50% ethanol, **3)** 50 mM TBS pH 7.5, **4)** 5x PBS pH 7.4, **5)** 50 mM MES and 180 mM NaCl pH 5.2, **6)** 100 mM $NaHCO_3$ pH 9.5, **7)** 500 mM pyridine in 50 mM TBS pH 7.5. Wells in a black 96-well plate (Grenier) were coated with 10 mM DOPA-Tet and incubated with 0.25 mM ALP (approximately 40 mg/mL) for 1 hour. The ALP solution was removed, and each well was washed twice with 300 µL of the chosen wash solution. For the "fast" group, a third wash was performed with the wash solution. For the "overnight" group, a third quantity of wash solution was added to each well and incubated for approximately 12 hours before being removed.

After washing, 100 µL 1 mg/mL p-NPP was added to each well and the change in absorbance at 405 nm was measured using the microplate reader.

Higher activity was observed in tubes washed with water and especially 50% ethanol (**Figure S7**). This suggests that these solutions were poor eluents, and the ionic buffers were better. No significant difference was observed between buffers of different pH, nor in the solvent with pyridine.

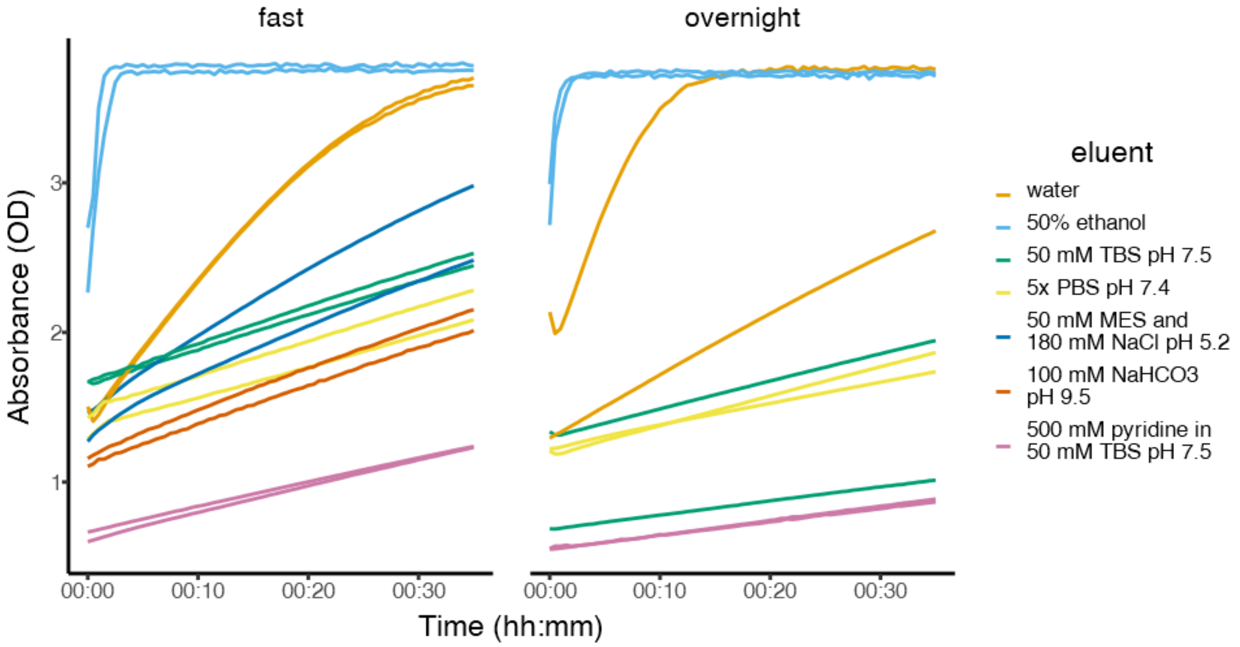


**Figure S7.** Residual ALP activity in coated/grafted microcentrifuge tubes incubated with 1 mg/mL p-NPP after washing with various wash solutions.

### Stability of Grafted ALP in Human Serum

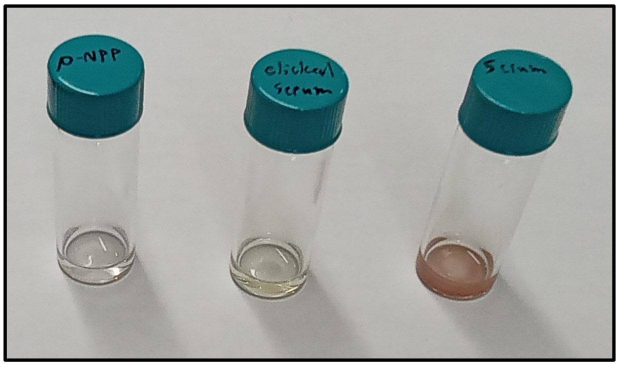


**Figure S8.** Effect of prolonged exposure to human serum on grafting. (Left) 1 mg/mL p-NPP, added for color comparison. (Middle) 1 mg/mL p-NPP incubated for 1 hour in serum-incubated, ALP-grafted vial. (Right) 1 mg/mL p-NPP incubated for 1 hour in serum-incubated, DOPA-Tet-coated vial.

### MTT Assay

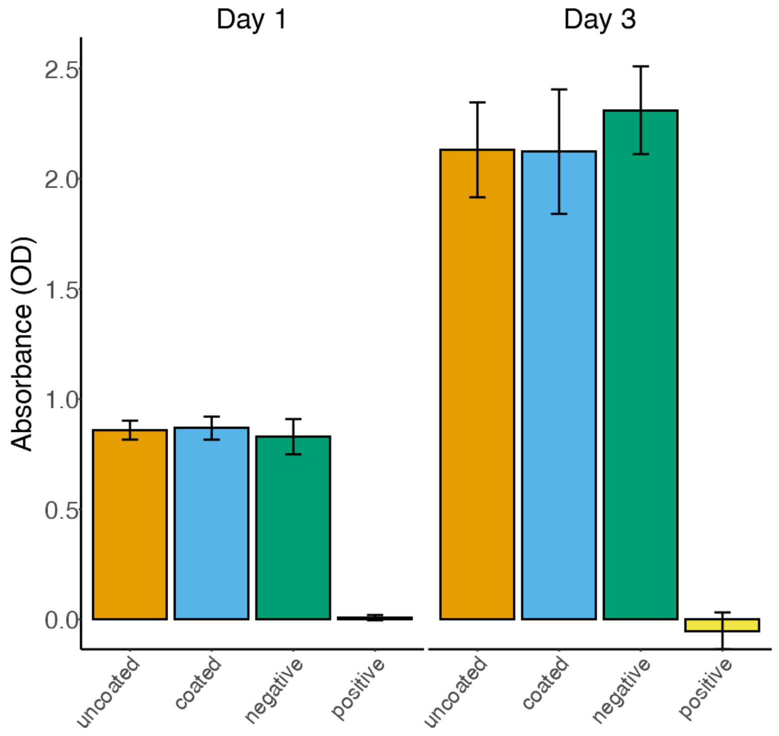


**Figure S9.** MTT Assay of extracts from uncoated and coated/(c(RGDfK)-grafted NanoECM after 1 day and 3 days culture with fibroblasts (NIH3T3). Error bars denote 1 standard deviation.

### Vancomycin Surface Inhibition of *S. aureus* Growth

**A**

| **MIC against *S. aureus*** | |
|---|---|
| Vancomycin | Vancomycin-TCO |
| 1 µg/mL | 16 µg/mL |

**B**

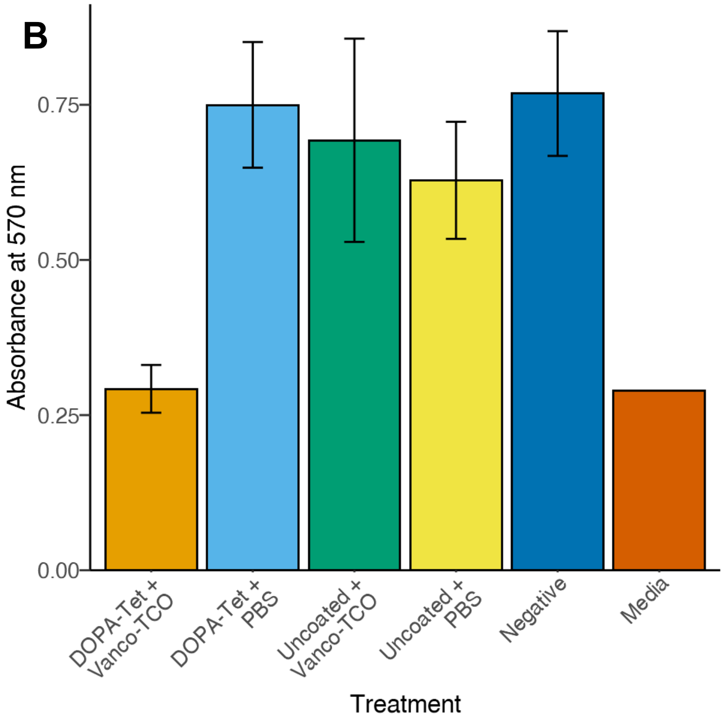


**Figure S10.** Inhibitory effect against *S. aureus* of both **(A)** in-solution and **(B)** surface-grafted vancomycin-TCO. **(A)** Minimum inhibitory concentration (MIC) of vancomycin and vancomycin-TCO against *S*. *aureus*. **(B)** Mean absorbance at 570 nm of *S. aureus* stained with PrestoBlue™, a resazurin-based cell viability indicator. Error bars denote 1 standard deviation.

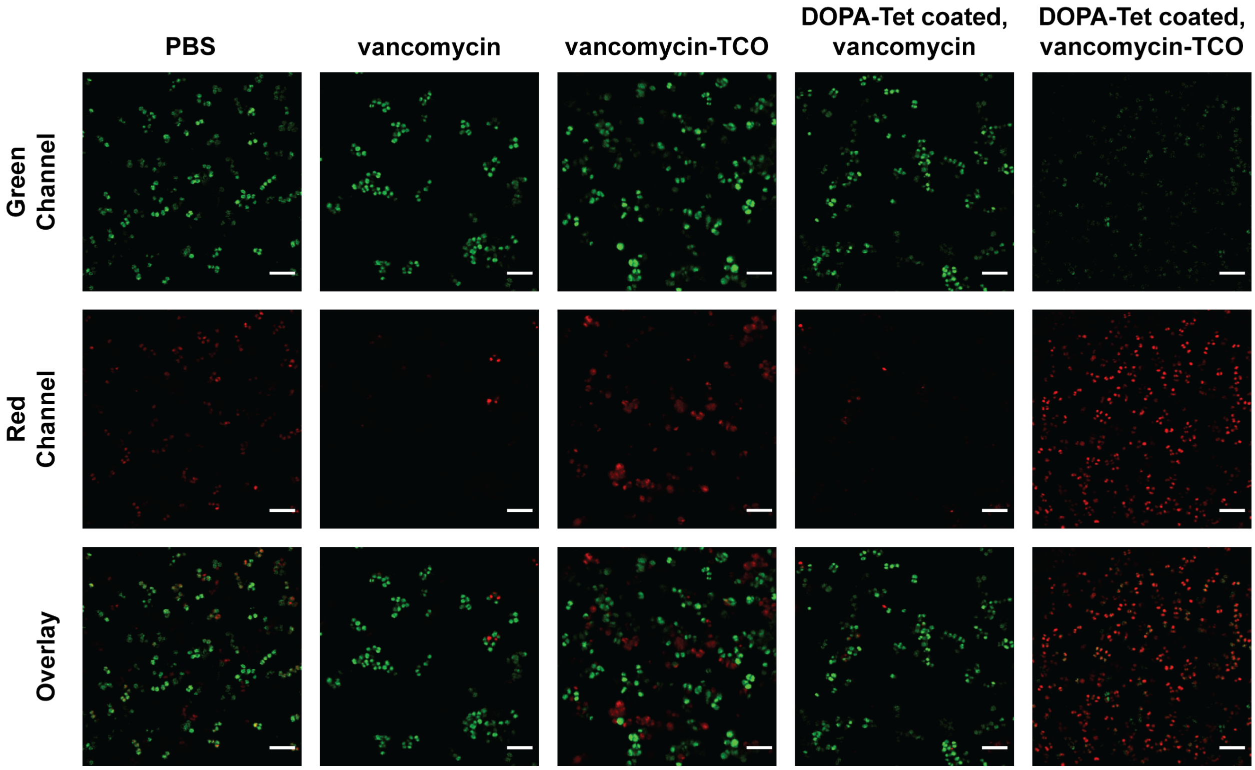


**Figure S11.** CLSM images of *S. aureus* culture incubated on surfaces grafted with vancomycin-TCO. Green channel: 488 nm, Red channel: 561 nm, Overlay: Overlay of the images acquired at the green and red channels. Cultures were stained with a combination of PI and SYTO 9, which selectively label dead (red) and live (green) cells, respectively. Scale bar is 5 μm.

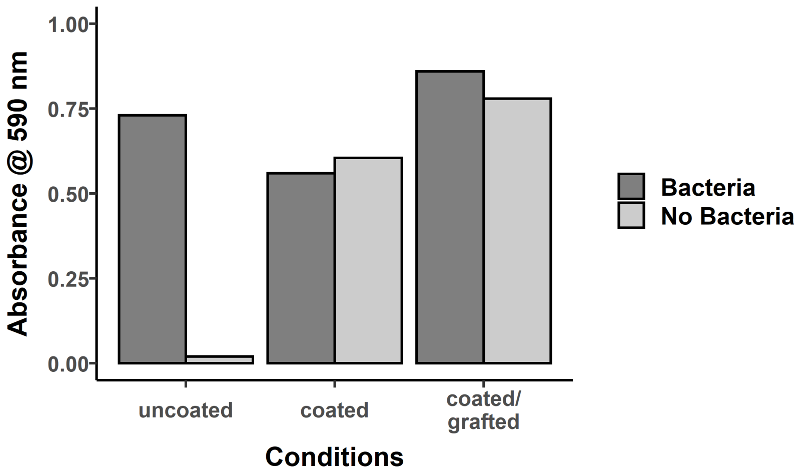


**Figure S12.** Absorbance values for crystal violet assay of uncoated, coated, and coated/vancomycin-grafted culture plates. Values shown are raw values, including the experimental data (bacteria) and negative controls (no bacteria) included. Median values are shown.
